# Bridging neuronal correlations and dimensionality reduction

**DOI:** 10.1101/2020.12.04.383604

**Authors:** Akash Umakantha, Rudina Morina, Benjamin R. Cowley, Adam C. Snyder, Matthew A. Smith, Byron M. Yu

**Affiliations:** Carnegie Mellon Neuroscience Institute, Pittsburgh, PA; Machine Learning Department, Carnegie Mellon University, Pittsburgh, PA; Department of Electrical and Computer Engineering, Carnegie Mellon University, Pittsburgh, PA; Princeton Neuroscience Institute, Princeton University, Princeton, NJ; Department of Brain and Cognitive Sciences, University of Rochester, Rochester, NY; Department of Neuroscience, University of Rochester, Rochester, NY; Center for Visual Science, University of Rochester, Rochester, NY; Department of Biomedical Engineering, Carnegie Mellon University, Pittsburgh, PA

## Abstract

Two commonly used approaches to study interactions among neurons are spike count correlation, which describes pairs of neurons, and dimensionality reduction, applied to a population of neurons. While both approaches have been used to study trial-to-trial correlated neuronal variability, they are often used in isolation and have not been directly related. We first established concrete mathematical and empirical relationships between pairwise correlation and metrics of population-wide covariability based on dimensionality reduction. Applying these insights to macaque V4 population recordings, we found that the previously reported decrease in mean pairwise correlation associated with attention stemmed from three distinct changes in population-wide covariability. Overall, our work builds the intuition and formalism to bridge between pairwise correlation and population-wide covariability and presents a cautionary tale about the inferences one can make about population activity by using a single statistic, whether it be mean pairwise correlation or dimensionality.

## Introduction

A neuron can respond differently to repeated presentations of the same stimulus. These variable responses are often correlated across pairs of neurons from trial to trial, measured using spike count correlations (*r*_sc_, also referred to as noise correlation; Cohen and Kohn, 2011). Studies have reported changes in spike count correlation across various experimental manipulations and cognitive phenomena, including attention (Cohen and Maunsell, 2009; Mitchell et al., 2009; Herrero et al., 2013; Gregoriou et al., 2014; Ruff and Cohen, 2014b; Snyder et al., 2018), learning (Gu et al., 2011; Jeanne et al., 2013; Ni et al., 2018), task difficulty (Ruff and Cohen, 2014a), locomotion (Erisken et al., 2014), stimulus drive (Maynard et al., 1999; Kohn and Smith, 2005; Smith and Kohn, 2008; Miura et al., 2012; Ponce-Alvarez et al., 2013; Ruff and Cohen, 2016b), decision-making (Nienborg et al., 2012), task context (Bondy et al., 2018), anesthesia (Ecker et al., 2010), adaptation (Adibi et al., 2013), and more (Fig. 1*a*). Spike count correlation also depends on timescales of activity (Bair et al., 2001; Kohn and Smith, 2005; Smith and Kohn, 2008; Mitchell et al., 2009; Runyan et al., 2017), neuromodulation (Herrero et al., 2013; Minces et al., 2017), and properties of the neurons themselves, including their physical distance from one another (Lee et al., 1998; Smith and Kohn, 2008; Smith and Sommer, 2013; Ecker et al., 2014; Solomon et al., 2015; Rosenbaum et al., 2017), tuning preferences (Lee et al., 1998; Romo et al., 2003; Kohn and Smith, 2005; Huang and Lisberger, 2009), and neuron type (Qi and Constantinidis, 2012; Snyder et al., 2016). Theoretical work has posited that changes in correlations affect neuronal computations and sensory information coding (Zohary et al., 1994; Shadlen and Newsome, 1998; Abbott and Dayan, 1999; Averbeck et al., 2006; Moreno-Bote et al., 2014; Sharpee and Berkowitz, 2019; Rumyantsev et al., 2020; Bartolo et al., 2020). Given such widespread empirical observations and theoretical insight, spike count correlation has been and remains instrumental in our current understanding of how neurons interact.

**Figure 1:**
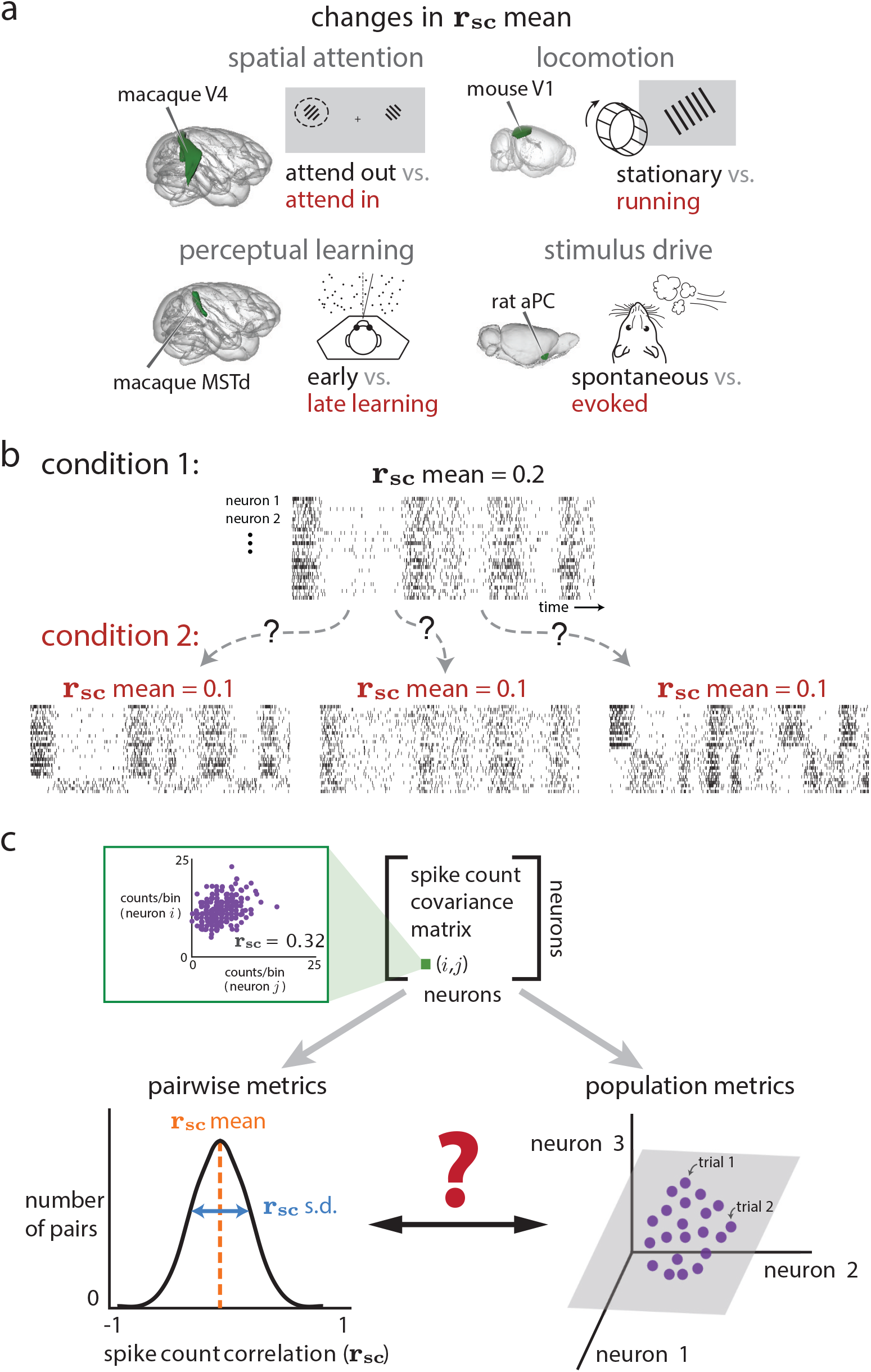
How do spike count correlations between pairs of neurons (i.e., pairwise metrics) relate to how the entire population co-fluctuates (i.e., population metrics)? **a.** Four example experiments in which mean spike count correlation (*r*_sc_ mean) has been observed to change between experimental conditions. These include spatial attention (macaque visual area V4; Cohen and Maunsell, 2009; Mitchell et al., 2009; Gregoriou et al., 2014; Luo and Maunsell, 2015; Snyder et al., 2018), perceptual learning (macaque dorsal medial superior temporal area; Gu et al., 2011), locomotion (mouse visual area V1; Erisken et al., 2014), and stimulus drive (rat anterior piriform cortex; Miura et al., 2012). **b.** The same change in *r*_sc_ mean (from 0.2 to 0.1 between conditions 1 and 2) could correspond to multiple distinct changes in the activity of the population of neurons. Condition 2, left: a decrease in *r*_sc_ mean could correspond to some neurons becoming anti-correlated with others in the population; in this case, some neurons that were previously positively correlated are now anti-correlated with the rest of the population (bottom rows of raster plot). Condition 2, middle: a decrease in *r*_sc_ mean could correspond to a decrease in how strongly neurons co-fluctuate together; in this case, neurons covary as in condition 1 but each neuron does not co-fluctuate with other neurons as strongly. Condition 2, right: a decrease in *r*_sc_ mean could correspond to the introduction of another ‘mode’ of covariation (i.e., an increase in the dimensionality of population activity); in this case, neurons in the top half of the raster covary as in condition 1, but neurons in the bottom half of the raster covary in a manner independent from those in the top half. **c**. Pairwise (*r*_sc_) and population (dimensionality reduction) metrics both arise from the same spike count covariance matrix, but the precise relationship between these two sets of metrics remains unknown. Top row: Each element of the spike count covariance matrix corresponds to the covariance across responses to repeated presentations of the same stimulus for two simultaneously-recorded neurons (e.g., neurons *i* and *j*, left inset). Bottom row: Pairwise metrics (left) typically summarize the distribution of spike count correlation with the mean (*r*_sc_ mean); in this work, we propose additionally reporting the standard deviation (*r*_sc_ s.d.). Population metrics (right) of the spike count covariance matrix are identified by applying dimensionality reduction to the population activity (e.g., gray plane depicts a low-dimensional space describing how neurons covary). By understanding the relationship between pairwise and population metrics, we can better interpret how changes in pairwise statistics (e.g., experiments in **a**) correspond to changes in population metrics, and vice-versa.

Most studies compute the average spike count correlation over pairs of recorded neurons for different experimental conditions, periods of time, neuron types, etc. A decrease in this mean correlation is commonly attributed to a reduction in the size (or gain) of shared co-fluctuations (Shadlen and Newsome, 1998; Rabinowitz et al., 2015; Lin et al., 2015; Ecker et al., 2016; Huang et al., 2019; Ruff et al., 2019b), e.g., a decrease in the strength of “common input” that drives each neuron in the population. However, other distinct changes at the level of the entire neuronal population can manifest as the same decrease in mean pairwise correlation (Fig. 1*b*). For example, a common input that drives the activity of all neurons up and down together could be altered to drive some neurons up and other neurons down. Alternatively, that first common input signal might remain the same, but a second input could be introduced that drives some neurons up and others down. It is difficult to differentiate these distinct possibilities using a single summary statistic, such as mean spike count correlation.

Distinguishing among these changes to the population-wide covariability might be possible by considering additional statistics that measure how the entire population of neurons co-fluctuates together. In particular, one may use dimensionality reduction to compute statistics that characterize multiple distinct features of population-wide covariability (Cunningham and Yu, 2014). Dimen-sionality reduction has been used to investigate decision-making (Harvey et al., 2012; Mante et al., 2013; Kiani et al., 2014; Kaufman et al., 2015), motor control (Churchland et al., 2012; Gallego et al., 2017), learning (Sadtler et al., 2014; Ni et al., 2018; Vyas et al., 2018), sensory coding (Mazor and Laurent, 2005; Pang et al., 2016), spatial attention (Cohen and Maunsell, 2010; Rabinowitz et al., 2015; Snyder et al., 2018; Huang et al., 2019), interactions between brain areas (Perich et al., 2018; Ruff and Cohen, 2019a; Ames and Churchland, 2019; Semedo et al., 2019; Veuthey et al., 2020), and network models (Williamson et al., 2016; Mazzucato et al., 2016; Recanatesi et al., 2019), among others. As with mean spike count correlation, the statistics computed from dimensionality reduction can also change with attention (Rabinowitz et al., 2015; Huang et al., 2019), stimulus drive (Churchland et al., 2010; Cowley et al., 2016; Snyder et al., 2018), motor output (Gallego et al., 2018), learning (Athalye et al., 2017), and anesthesia (Ecker et al., 2014). However, unlike mean spike count correlation (henceforth referred to as a “pairwise metric”) which averages across pairs of neurons, the statistics computed from dimensionality reduction (henceforth referred to as “population metrics”) consider the structure of population-wide covariability (Fig. 1*c*). An example of a commonly reported population metric is dimensionality (Yu et al., 2009; Rabinowitz et al., 2015; Cowley et al., 2016; Williamson et al., 2016; Mazzucato et al., 2016; Gao and Ganguli, 2015; Gallego et al., 2017; Stringer et al., 2019a; Recanatesi et al., 2019). Dimensionality is used to assess whether the number of population co-fluctuation patterns (possibly reflecting the number of common inputs) changes across experimental conditions (Fig. 1*b*, condition 1 versus condition 2, right panel). Thus, population metrics could help to distinguish among the distinct ways in which population-wide covariability can change, especially those that lead to the same change in mean spike count correlation (Fig. 1*b*).

Both pairwise and population metrics aim to characterize how neurons covary, and both can be computed from the *same* spike count covariance matrix (Fig. 1*c*). Still, studies rarely report both, and the relationship between the two is not known. In this study, we establish the relationship between pairwise metrics and population metrics both analytically and empirically using simulations. We find that changes in mean spike count correlation could correspond to several distinct changes in population metrics including: 1) the strength of shared variability (e.g., the strength of a common input), 2) whether neurons co-fluctuate together or in opposition (e.g., how similarly a common input drives each neuron in the population), or 3) the dimensionality (e.g., the number of common inputs). Furthermore, we show that a rarely-reported statistic–the standard deviation of spike count correlation–provides complementary information to the mean spike count correlation about how a population of neurons co-fluctuates. Applying this understanding to recordings in area V4 of macaque visual cortex, we found that the previously-reported decrease in mean spike count correlation with attention stems from multiple distinct changes in population-wide covariability. Overall, our results demonstrate that common ground exists between the literatures of spike count correlation and dimensionality reduction and provides a cautionary tale for attempting to draw conclusions about how a population of neurons covaries using one, or a small number of, statistics. Our framework builds the intuition and formalism to navigate between the two approaches, allowing for a more interpretable and richer description of the interactions among neurons.

## Results

### Defining pairwise and population metrics

We first define the metrics that we will use to summarize 1) the distribution of spike count correlations (i.e., pairwise metrics) and 2) dimensionality reduction of a population covariance matrix (i.e., population metrics). For pairwise metrics, we consider the mean and standard deviation (s.d.) of *r*_sc_ across all pairs of neurons, which summarize the *r*_sc_ distribution (Fig. 1*c*, bottom left panel). For population metrics, we consider loading similarity, percent shared variance (abbreviated to %sv), and dimensionality (described below and in more detail in Methods). These metrics each describe some aspect of population-wide covariability and thus represent natural, multivariate extensions of *r*_sc_.

To illustrate these three population metrics, consider the activity of a population of neurons over time (Fig. 2*a*, spike rasters). If the activity of all neurons goes up and down together, we would find the pairwise spike count correlations between all pairs of neurons to be positive. A more succinct way to characterize this population activity is to identify a single time-varying *latent co-fluctuation* that is shared by all neurons (Fig. 2*a*, blue line). The way in which neurons are coupled to this latent co-fluctuation is indicated by a *loading* for each neuron. In this example, because the latent co-fluctuation describes each neuron’s activity going up and down together, the loadings have the same sign (Fig. 2*a*, green rectangles). We refer to the latent co-fluctuation’s corresponding set of loadings as a *co-fluctuation pattern*. A co-fluctuation pattern can be represented as a direction in the population activity space, where each coordinate axis corresponds to the activity of one neuron (Fig. 2*a*, right panel).

**Figure 2:**
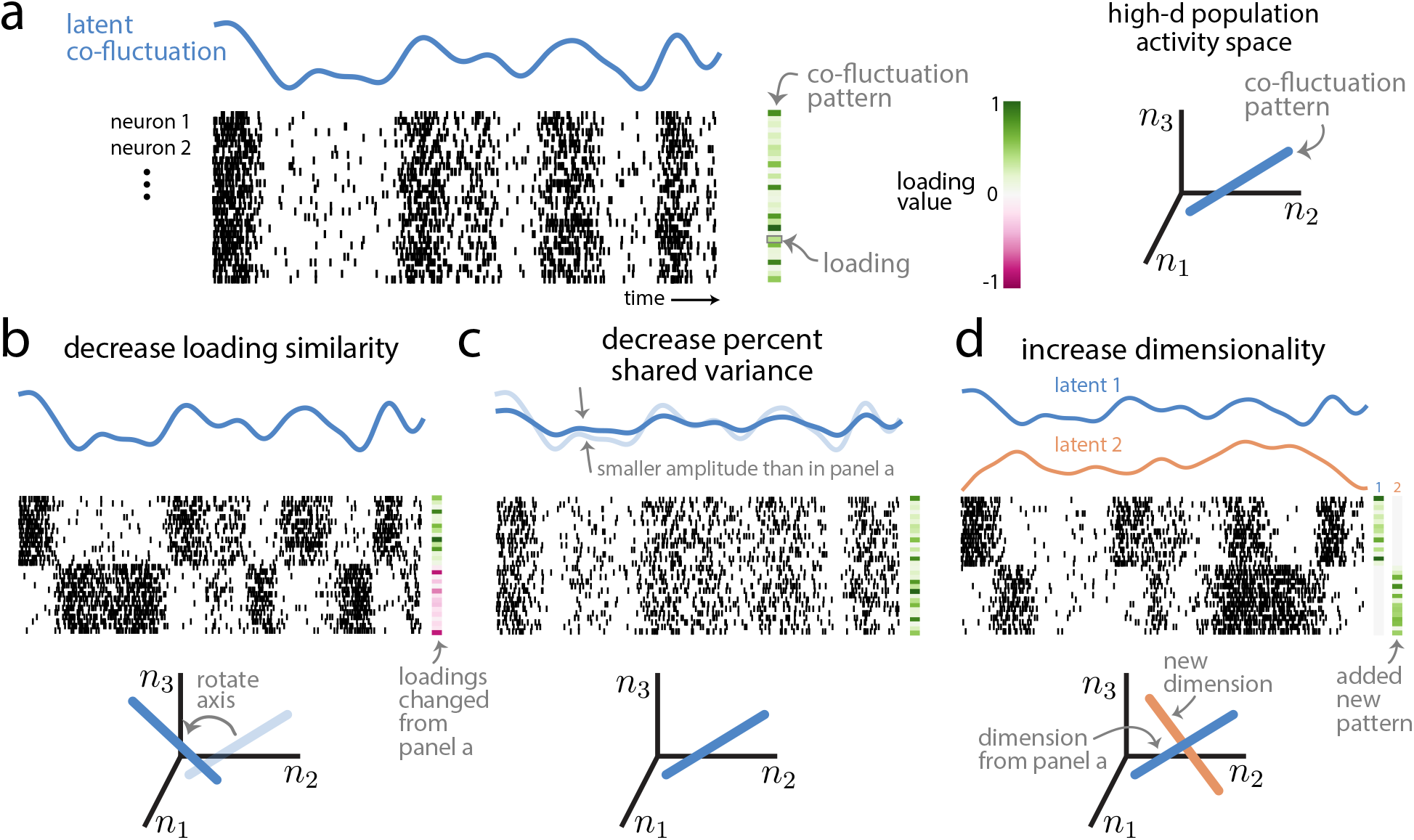
Intuition about population metrics: loading similarity, percent shared variance (%sv), and dimensionality. **a**. Population activity (population raster, where each row is the spike train for one neuron over time) is characterized by a latent co-fluctuation (blue) and a co-fluctuation pattern made up of loadings (green rectangles). Each neuron’s underlying firing rate is a product of the latent and that neuron’s loading (which may either be positive or negative). One may also view population activity through the lens of the population activity space (right plot), where each axis represents the activity of one neuron (*n*_1_, *n*_2_, *n*_3_ represent neuron 1, neuron 2, and neuron 3). In this space, a co-fluctuation pattern corresponds to an axis whose orientation depends on the pattern’s loadings (right plot, blue line). **b**. Population activity with a lower loading similarity than in panel **a**. The loadings have both positive and negative values (i.e., dissimilar loadings), leading to neurons that are anti-correlated (compare top rows with bottom rows of population raster). Changing the loading similarity will rotate a pattern’s axis in the population activity space (bottom plot, ‘rotate axis’). **c**. Population activity with a lower %sv than in panel **a**. The latent co-fluctuation shows smaller amplitude changes over time than in panel **a**, which leads to a lower %sv. Changing %sv leads to no changes of the co-fluctuation pattern (bottom plot, axis is same as that in panel **a**). **d**. Population activity with a dimensionality of 2, compared to a dimensinality of 1 in panel **a**. Adding a new dimension leads to a new latent co-fluctuation (orange line) and a new co-fluctuation pattern (‘new dimension’). Each neuron’s underlying firing rate is expressed as a weighted combination of the latents, where the weights correspond the neuron’s loadings in each co-fluctuation pattern. Here, each dimension corresponds to a distinct subset of neurons (top rows vs. bottom rows); in general, this need not be the case, as each neuron typically has nonzero weights for both dimensions. In the population activity space (bottom plot), the activity varies along the two axes (i.e., a 2-d plane) defined by the two co-fluctuation patterns.

The first population metric is *loading similarity*, a value between 0 and 1 that describes to what extent the loadings differ across neurons within a co-fluctuation pattern. A loading similarity close to 1 indicates that the loadings have the same sign and are of similar magnitude (Fig. 2*a*, green rectangles). A loading similarity close to 0 indicates that many of the loadings differ, either in magnitude, sign, or both (Fig. 2*b*, green and pink squares). In this case, some neurons may have positive loadings and co-fluctuate in the same direction as the latent co-fluctuation (Fig. 2*b*, top rows of neurons show high firing rates when blue line is high and low firing rates when blue line is low), whereas other neurons may have negative loadings and co-fluctuate in the opposite direction as the latent co-fluctuation (Fig. 2*b*, bottom rows of neurons show low firing rates when blue line is high and high firing rates when blue line is low). One can view changing the loading similarity as rotating the direction of a co-fluctuation pattern in population activity space (Fig. 2*b*, bottom plot).

The second population metric is *percent shared variance* or %sv, which measures the percentage of spike count variance explained by the latent co-fluctuation. This percentage is computed per neuron, then averaged across all neurons in the population (Williamson et al., 2016). A %sv close to 100% indicates that the activity of each neuron is tightly coupled to the latent co-fluctuation, with a small portion of variance that is independent to each neuron (Fig. 2*a*). A %sv close to 0% indicates that neurons fluctuate almost independently of each other and their activity weakly adheres to the time course of the latent co-fluctuation (Fig. 2*c*). By changing %sv, one does not change the co-fluctuation pattern in population activity space (Fig. 2, blue lines are the same in panels *a* and *c*) but rather the strength of the latent co-fluctuation (Fig. 2*c*, blue line has smaller amplitude than in panel *a*).

The third population metric is *dimensionality*. We define dimensionality as the number of co-fluctuation patterns (or dimensions) needed to explain the shared variability among neurons (see Methods). The variable activity of neurons may depend on multiple common inputs, e.g., top-down signals like attention and arousal (Rabinowitz et al., 2015; Cowley et al., 2020) or spontaneous and uninstructed behaviors (Stringer et al., 2019b; Musall et al., 2019). Furthermore, these common inputs may differ in how they modulate neurons. This may result in two or more dimensions of the population activity (Fig. 2*d*, blue and orange latent co-fluctuations). For illustrative purposes, each dimension might correspond to a single group of tightly-coupled neurons (Fig. 2*d*, neurons in top rows have non-zero loadings for pattern 1, whereas neurons in bottom rows have non-zero loadings for pattern 2). However, in general, each neuron can have non-zero loadings for multiple patterns. In population activity space, adding a new dimension adds a new axis along which neurons covary (Fig. 2*d*, orange line). We use the term *dimension* to refer either to a latent co-fluctuation or its corresponding co-fluctuation pattern, depending on context.

### Varying population metrics to assess changes in pairwise metrics

Given that both pairwise and population metrics are computed from the same spike count covariance matrix (Fig. 1*c*), a connection should exist between the two. We establish this connection by deriving mathematical relationships and carrying out simulations. In simulations, we assessed how systematically changing one of the population metrics (e.g., increasing loading similarity, Fig. 3*a*), changes the spike count covariance matrix (Fig. 3*b*), and the corresponding *r*_sc_ distribution (Fig. 3*c*), which we summarized using its mean and standard deviation (Fig. 3*d*). The covariance matrix was parameterized in a way that allowed us to create covariance matrices with specified population metrics (see Methods). Thus, our simulation procedure does not simulate neuronal activity, but rather creates covariance matrices which are consistent with the specified population metrics.

**Figure 3:**
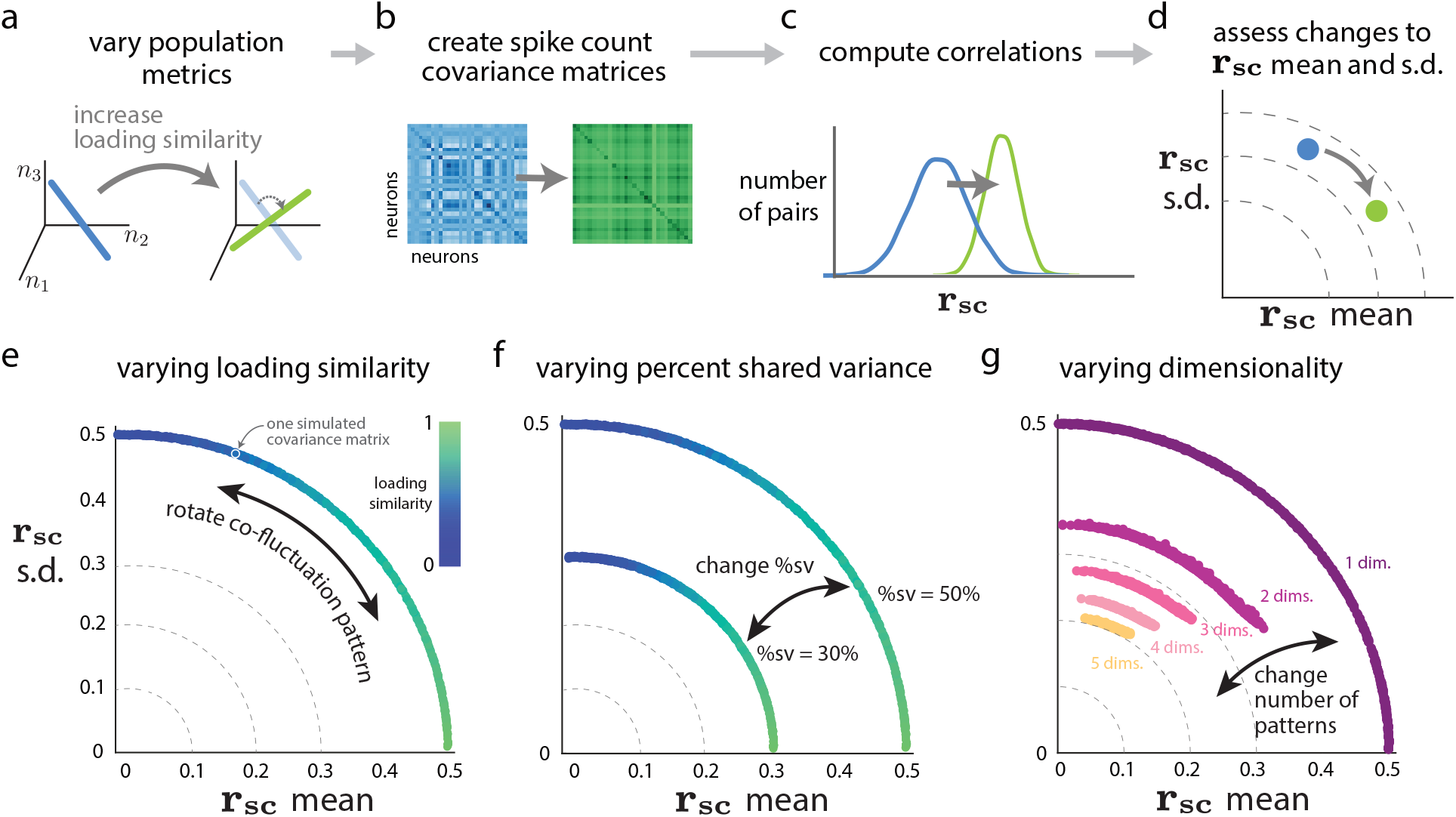
Relationship between population metrics and pairwise metrics. Panels **a**-**d** describe the simulation procedure to assess how systematic changes in population metrics lead to changes in pairwise metrics. **a**. We first systematically varied one of the population metrics while keeping the others fixed. For example, we can increase the loading similarity from a low value (left, blue) to a high value (right, green), while keeping %sv and dimensionality fixed. **b**. Then, we constructed covariance matrices corresponding to each value of the population metric in panel **a** (see Methods), without generating synthetic data. **c**. For each covariance matrix from panel **b**, we directly computed the correlations (i.e., the *r*_sc_ distributions). **d**. We computed *r*_sc_ mean and *r*_sc_ s.d. from the *r*_sc_ distributions in panel **c** and then assessed how the change in a given population metric from panel **a** changed pairwise metrics. In this case, the increase in loading similarity increased *r*_sc_ mean and decreased *r*_sc_ s.d. (blue dot to green dot). **e**. Varying loading similarity with a fixed %sv of 50% and dimensionality of 1. Each dot corresponds to the *r*_sc_ mean and *r*_sc_ s.d. of one simulated covariance matrix with specified population metrics (dots are close together and appear to form a continuum). The color of each dot corresponds to the loading similarity (see Methods), where a value of 1 indicates that all loading weights have the same value. **f**. Varying %sv. The same setting as in panel **e**, except we consider two different values of percent shared variance (50% and 30%). **g**. Varying dimensionality (i.e., number of co-fluctuation patterns) while sweeping loading similarity between 0 and 1 and keeping %sv fixed at 50%. In this simulation, the relative strengths of each dimension uniform across dimensions (i.e., flat eigenspectra; see Methods).

#### Loading similarity has opposing effects on rsc mean and s.d

We first asked how the loading similarity of a single co-fluctuation pattern (i.e., one dimension) affected *r*_sc_ mean and s.d. Intuitively, a high loading similarity indicates that the activity of all neurons increases and decreases together (Fig. 2*a*), resulting in values of *r*_sc_ that are all positive and similar in value. Indeed, in simulations, we found that high loading similarity corresponded to large *r*_sc_ mean and *r*_sc_ s.d. close to 0 (Fig. 3*e*, green dots near horizontal axis). On the other hand, a low loading similarity indicates that when some neurons increase their activity, others decrease their activity (Fig. 2*b*), resulting in some positive *r*_sc_ values (for pairs that change their activity in the same direction) and some negative *r*_sc_ values (for pairs that change their activity in opposition). In simulations, a low loading similarity indeed corresponded to an *r*_sc_ mean close to 0 and a large *r*_sc_ s.d. (Fig. 3*e*, blue dots near vertical axis). By varying the loading similarity, we surprisingly observed an arc-like trend in the *r*_sc_ mean versus *r*_sc_ s.d. plot (Fig. 3*e*). In Supplementary Math Note A, we derive the analytical relationship between loading similarity and *r*_sc_. In Supplementary Math Note B, we show mathematically why the *r*_sc_ mean versus *r*_sc_ s.d. relationship follows a circular arc.

#### Decreasing %sv reduces r_*sc*_ mean and s.d

We next asked how %sv, which measures the percentage of each neuron’s variance that is shared with other neurons in the population, is related to *r*_sc_ mean and s.d. Intuitively, one might expect %sv and *r*_sc_ mean to be closely related because *r*_sc_ measures the degree to which the activity of two neurons is shared (Cohen and Kohn, 2011). We investigated this in simulations and found that how closely %sv and *r*_sc_ mean were related depended on the loading similarity. When loading similarity was high (Fig. 3*f*, green dots), there was a direct relationship between %sv and *r*_sc_ mean (specifically, %sv equals *r*_sc_ mean). However, when loading similarity was low (Fig. 3*f*, blue dots), the relationship between %sv and *r*_sc_ mean was less direct. Namely, *r*_sc_ mean remained close to zero regardless of %sv. This illustrates that *r*_sc_ mean and %sv are not the same. It is possible for a population of neurons with high %sv (e.g., Fig. 3*f*, blue dots in outer arc) to have smaller *r*_sc_ mean than a population with lower %sv (e.g., Fig. 3*f*, green dots in inner arc).

These relationships that we have shown through simulation can be captured mathematically. First, if we have knowledge of the loading weights in the co-fluctuation pattern, the *r*_sc_ between a pair of neurons can be expressed in terms of the %sv and loading values of the two neurons (Supplementary Math Note A):

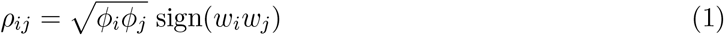

where *ρ*_*ij*_ is the *r*_sc_ between neurons *i* and *j*, *ϕ*_*i*_ and *ϕ*_*j*_ are the %sv of each neuron (expressed as a proportion per neuron, in contrast to %sv in Fig. 3*f* which shows the average %sv across all neurons), and *w*_*i*_ and *w*_*j*_ are the loadings of the neurons in the co-fluctuation pattern. The *r*_sc_ mean is the average of *ρ*_*ij*_ values across all neuron pairs. From equation (1), we observe that when loading similarity is high (i. most loading weights have the same sign), %sv and *r*_sc_ mean are directly related (i.e., 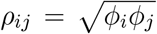). However, when loading similarity is low (i.e., some loading weights are positive and others are negative), *r*_sc_ mean is small regardless of %sv because some pairs have sign(*w*_*i*_*w*_*j*_) = +1 and others have sign(*w*_*i*_*w*_*j*_) = −1.

Second, if we have information about the *r*_sc_ s.d. (instead of loading weights), we can establish the following relationship between %sv, *r*_sc_ mean, and *r*_sc_ s.d. (Supplementary Math Note B):

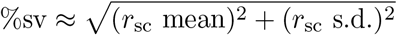

In other words, in the *r*_sc_ mean versus *r*_sc_ s.d. plot, %sv is reflected in the distance of a point from the origin (Fig. 3*f*). This relationship holds regardless of the loading similarity. The intuition is that the %sv corresponds to the magnitude of *r*_sc_ values (i.e., the |*ρ*_*ij*_| from equation (1)).

These findings highlight the pitfalls of considering a single statistic (e.g., *r*_sc_ mean) on its own and the benefits of considering multiple statistics (e.g., both *r*_sc_ mean and s.d.) when trying to draw conclusions about how neurons covary. By considering *r*_sc_ mean and s.d. together, one can gain insight into the loading similarity (Fig. 3*e*) and the %sv (Fig. 3*f*) of a neuronal population. Thus far, we have only considered the specific case where activity co-fluctuates along a single dimension in the firing rate space. We next considered how pairwise metrics change in the more general case where neuronal activity co-fluctuates along multiple dimensions.

#### Adding more dimensions tends to reduce r_sc_ mean and s.d

We sought to assess how dimensionality (i.e., the number of co-fluctuation patterns) is related to pairwise metrics. In simulations, we increased the number of co-fluctuation patterns (compare Fig. 2*a* to *d*; see Methods), while sweeping loading similarity and fixing the total %sv. We found that increasing dimensionality tended to reduce *r*_sc_ mean and s.d. (Fig. 3*g*, dots for larger dimensionalities lay closer to the origin than dots for smaller dimensionalities).

It seems counterintuitive that adding a new way in which neurons covary reduces the magnitude of *r*_sc_. The intuition is that if multiple distinct (i.e., orthogonal) dimensions exist, then a neuron pair interacts in opposing ways along different dimensions. For example, consider two neurons with loadings of the same sign in one co-fluctuation pattern, and opposite sign in the second pattern. If only the first dimension exists, the two neurons would go up and down together and be positively correlated. If only the second dimension exists, the two neurons would co-fluctuate in opposition and be negatively correlated. When both dimensions exist, the positive correlation from the first dimension and the negative correlation from the second dimension offset, and the resulting correlation between the neurons would be smaller than if only the first dimension were present. We formalize the above intuition in Supplementary Math Note C. We also show analytically that increasing dimensionality tends to move points closer to the origin in the *r*_sc_ mean versus *r*_sc_ s.d. plot (i.e., decrease *r*_sc_ mean and s.d.; Supplementary Math Note D).

However, we note that an increase in dimensionality does not imply that *both r*_sc_ mean and *r*_sc_ s.d. necessarily decrease. For example, in the case where the first co-fluctuation pattern has high loading similarity, adding more dimensions means it is less likely for *r*_sc_ s.d. to be 0 (Fig. 3*g*, compare dot closest to horizontal axis for ‘1 dim.’ to that for ‘2 dims.’). The intuition is that if the first co-fluctuation pattern has a loading similarity of 1, the loading weights for all neurons are the same and thus *r*_sc_ values between all pairs are the same, resulting in *r*_sc_ s.d. of 0. Adding an orthogonal dimension to this pattern necessarily means adding a pattern with low loading similarity (Supplementary Math Note E), making it less likely for *r*_sc_ across all pairs to be the same. Therefore, *r*_sc_ s.d. is unlikely to be 0 for two dimensions (Fig. 3*g*, the smallest *r*_sc_ s.d. for ‘2 dims.’ is around 0.2). Still, in Figure 3*g* the dots for ‘2 dims.’ are closer to the origin than the dots for ‘1 dim’, implying that even if *r*_sc_ s.d. increases with an increase in dimensionality, the *r*_sc_ mean must decrease to a larger extent (Supplementary Math Note D).

#### The relative strength of each dimension impacts pairwise metrics

In the previous simulation (Fig. 3*g*), we assumed that each dimension explained an equal proportion of the overall shared variance (e.g., for two dimensions, each dimension explained half of the shared variance; see Methods). However, it is typically the case for recorded neuronal activity that some dimensions explain more shared variance than others; in other words, neuronal activity co-fluctuates more strongly along some patterns than others (Sadtler et al., 2014; Williamson et al., 2016; Mazzucato et al., 2016; Gallego et al., 2018; Huang et al., 2019; Stringer et al., 2019a; Ruff et al., 2019b). We sought to assess the influence of the relative strength of each dimension on pairwise metrics.

We reasoned that stronger dimensions would play a larger role than weaker dimensions in determining the *r*_sc_ distribution and pairwise metrics. Extending equation (1) to multiple dimensions, we show that the *r*_sc_ between a pair of neurons can be expressed as the sum of a contribution from each constituent dimension (Supplementary Math Note C). The stronger a dimension, the larger the magnitude of its contribution to *r*_sc_, and thus the larger its impact on *r*_sc_ mean and s.d.

To test this empirically, we performed a simulation with two dimensions, while systematically varying the relative strength of each dimension. We considered two scenarios: (1) one dimension has a pattern with high loading similarity and one dimension has a pattern with low loading similarity (Fig. 4*a*), and (2) both dimensions have patterns with low loading similarity (Fig. 4*b*). Note that both dimensions cannot have patterns with high loading similarity because they would not be orthogonal (Supplementary Math Note E).

**Figure 4:**
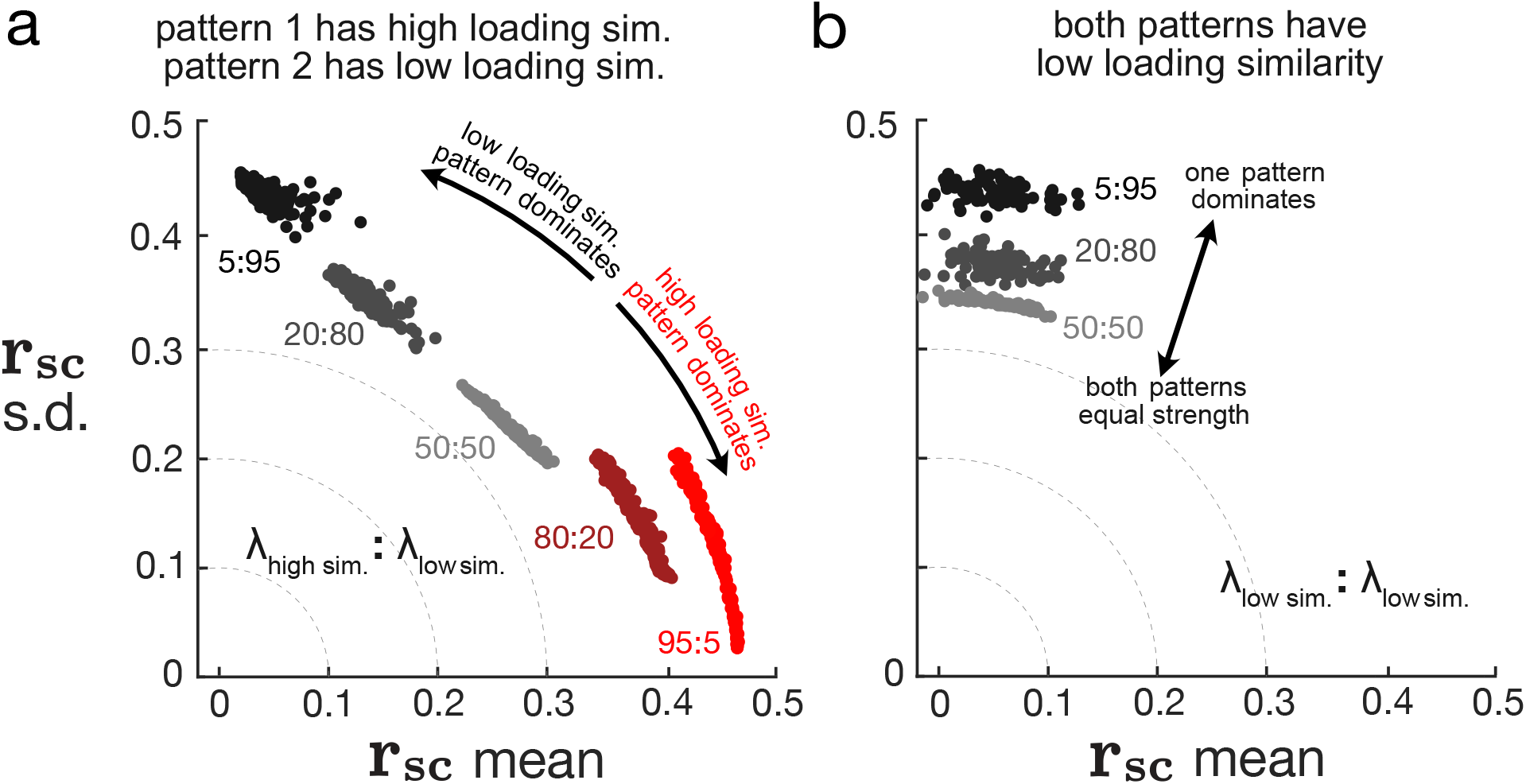
Relative strengths of dimensions affect *r*_sc_ distributions. With dimensionality of 2, we systematically varied the relative strengths of the two dimensions with a fixed total %sv of 50%. We considered two scenarios: 1) one dimension has high loading similarity and the other dimension has low loading similarity (panel **a**) and 2) both dimensions have low loading similarity (panel **b**). Each dot represents one simulated covariance matrix and *r*_sc_ distribution. The color of the dots indicate different relative strengths between the two dimensions, and numbers next to each cloud of dots indicate the ratio between the relative strength associated with each dimension. For example, in panel **a**, red dots correspond to the high loading similarity dimension being 19 times stronger (95:5) than the low loading similarity dimension. Black dots correspond to the low loading similarity dimension being 19 times stronger (5:95) than the high loading similarity dimension. In panel **b**, since both patterns have low loading similarity, clouds for 80:20 and 95:5 are very similar to clouds for 20:80 and 5:95 respectively and are thus omitted for clarity.

In scenario (1) where one dimension’s pattern has high loading similarity and the other has low loading similarity, *r*_sc_ mean and *r*_sc_ s.d. reflects the loading similarity of the dominant dimension (Fig. 4*a*). When the dimension with a high loading similarity pattern dominated, *r*_sc_ mean was large and *r*_sc_ s.d. was small (Fig. 4*a*, red dots are close to horizontal axis). When the dimension with a low loading similarity pattern dominated, *r*_sc_ mean was small and *r*_sc_ s.d. was large (Fig. 4*a*, black dots are close to vertical axis). When the two dimensions were of equal strength (i.e., neither dimension dominated), *r*_sc_ mean and *r*_sc_ s.d. were both intermediate values (Fig. 4*a*, light gray dots are between red and black dots). Thus, the dimensions along which neuronal activity co-fluctuates most strongly have a greater influence on pairwise metrics.

In scenario (2) where both dimensions have patterns of low loading similarity, *r*_sc_ mean was low and *r*_sc_ s.d. was high (Fig. 4*b*), similar to when there is one dimension with low loading similarity (Fig. 3*e*, blue dots). When we made one dimension stronger than the other, *r*_sc_ mean remained low and *r*_sc_ s.d. remained high (Fig. 4*b*, light gray dots and black dots are both close to vertical axis) because both patterns had low loading similarity. However, the radius of the arc increased (Fig. 4*b*, black dots farther from the origin than light gray dots), and was close to the arc that would have been produced with a single dimension (Fig. 3*g*, ‘1 dim.’). Thus, whereas changing the number of dimensions causes discrete jumps in the arc radius (Fig. 3*g*), changing the relative strength of each dimension allows for *r*_sc_ mean and *r*_sc_ s.d. to vary continuously between the arcs for different dimensionalities. Put another way, changing the relative strength of each dimension varies the “effective dimensionality” of population activity in a continuous manner. Neuronal activity for which one dimension dominates another (Fig. 4*b*, black dots) has a lower effective dimensionality than when both dimensions have equal strength (Fig. 4*b*, light gray dots).

### Reporting only a single statistic provides an incomplete description of population covariability

Figure 5 summarizes the relationships that we have established between pairwise metrics and population metrics. Rotating a co-fluctuation pattern from a low loading similarity to a high loading similarity increases *r*_sc_ mean and decreases *r*_sc_ s.d. along an arc (Fig. 5, arrow outside pink arc). Decreasing %sv decreases both *r*_sc_ mean and s.d. (Fig. 5, arrow pointing toward origin), and increasing dimensionality also tends to decrease *r*_sc_ mean and s.d. (Fig. 5, pink to yellow shaded regions).

**Figure 5:**
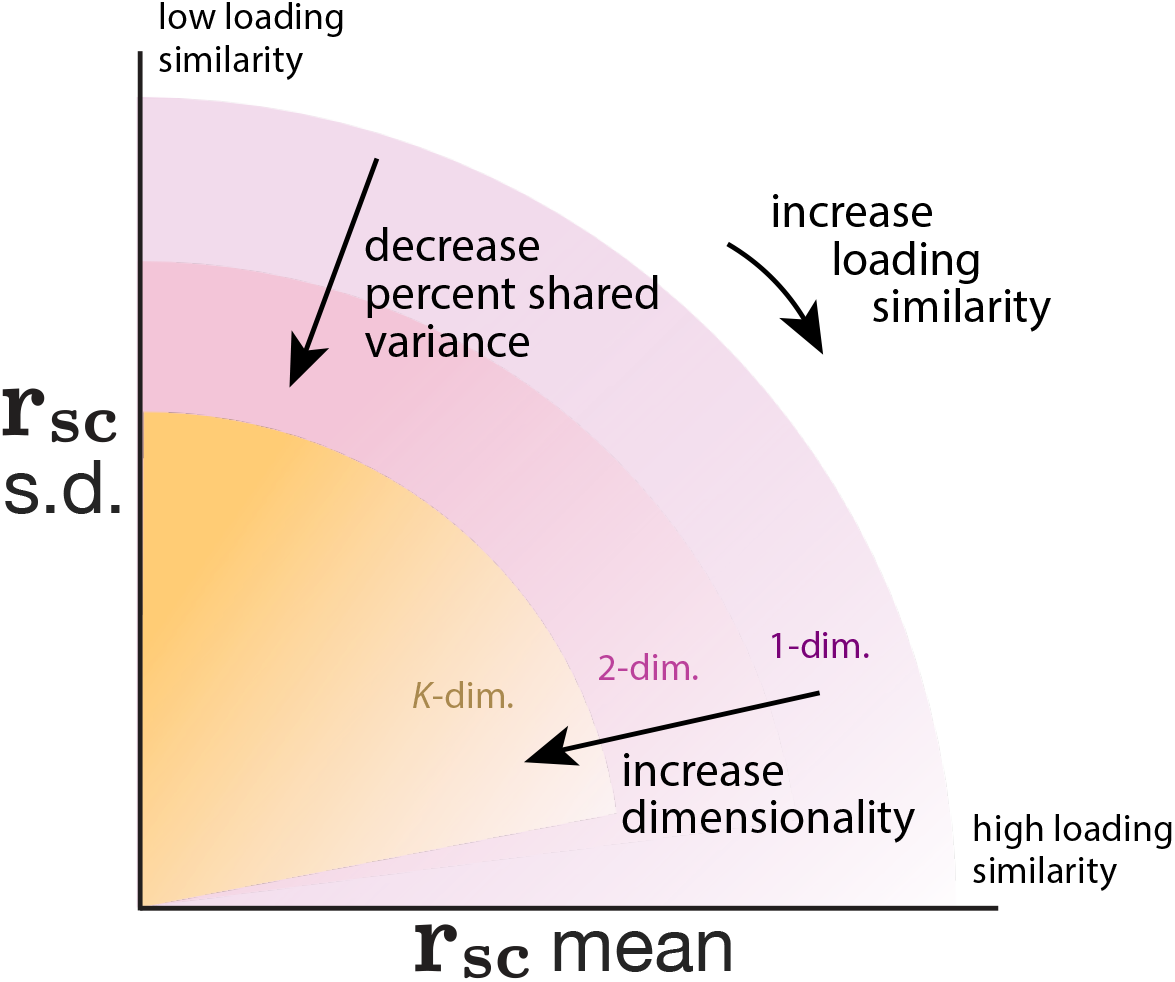
Summary of relationship between pairwise and population metrics. A change in *r*_sc_ mean and *r*_sc_ s.d. may correspond to changes in loading similarity, %sv, dimensionality, or a combination of the three. Shaded regions indicate the possible *r*_sc_ mean and *r*_sc_ s.d. values for different dimensionalities; increasing dimensionality tends to decrease *r*_sc_ mean and *r*_sc_ s.d. (shaded regions for larger dimensionalities become smaller). Within each shaded region, decreasing %sv decreases both *r*_sc_ mean and s.d. radially toward the origin. Finally, rotating co-fluctuation patterns such that the loadings are more similar (going from low to high loading similarity) results in moving clockwise along an arc such that *r*_sc_ mean increases and *r*_sc_ s.d. decreases. We also note two subtle trends. First, there are more possibilities for loading similarity to be low than high (Supplementary Math Note E), suggesting that *r*_sc_ s.d. will generally tend to be larger than *r*_sc_ mean if neuronal activity varied along a randomly chosen co-fluctuation pattern (shading within each region is darker near the vertical axis than the horizontal axis). Second, this effect becomes exaggerated for higher-dimensional neuronal activity as many dimensions can have low loading similarity but only one dimension can have high loading similarity (Supplementary Math Note E). Thus, it becomes progressively unlikely for *r*_sc_ s.d. to be 0 as dimensionality increases (shaded regions for larger dimensionalities lifted off the horizontal axis).

These results provide a cautionary tale that using a single statistic on its own provides an opaque description of population-wide covariability. For example, a change in *r*_sc_ mean could correspond to changes in loading similarity, %sv, dimensionality, or a combination of the three. Likewise, reporting dimensionality on its own would be incomplete because the role of a dimension in explaining population-wide covariability depends how much shared variance it explains and the loading similarity of its co-fluctuation pattern. For example, consider a decrease in dimensionality by 1. This would have little impact on population-wide covariability if the removed dimension explains only a small amount of shared variance, whereas it could have a large impact if the removed dimension explains a large amount of shared variance.

Considering multiple statistics together provides a richer description of population-wide covariability. For example, in the case where population activity co-fluctuates along a single dimension, *r*_sc_ mean and *r*_sc_ s.d. can be used together to approximate %sv (using distance from the origin) and deduce whether loading similarity is low (*r*_sc_ s.d. > *r*_sc_ mean) or high (*r*_sc_ mean > *r*_sc_ s.d.), whereas *r*_sc_ mean alone would not provide much information about %sv or loading similarity (cf. Fig. 5). In the next section, we further demonstrate using neuronal recordings how relating pairwise and population metrics using the framework we have developed (Fig. 5) provides a richer description of how neurons covary than using a single statistic (e.g., *r*_sc_ mean) alone.

### Case study: V4 neuronal recordings during spatial attention

When spatial attention is directed to the receptive fields of neurons in area V4 of macaque visual cortex, *r*_sc_ mean among those neurons decreases (Cohen and Maunsell, 2009; Mitchell et al., 2009; Gregoriou et al., 2014; Snyder et al., 2016, 2018). This decrease has often been attributed to a reduction in shared modulations among the neurons. However, we have shown both mathematically and in simulations that several distinct changes in population metrics (e.g., decrease in loading similarity, decrease in %sv, or an increase in dimensionality) could underlie this decrease in *r*_sc_ mean (Fig. 5). Here, we sought to assess which aspects of population-wide covariability underlie, and how each of them contribute to, the overall decrease in *r*_sc_ mean.

We analyzed activity recorded simultaneously from tens of neurons in macaque V4 while the animal performed an orientation-change detection task (Fig. 6*a*; previously reported in Snyder et al., 2018). To probe spatial attention, we cued the animal to the location of the stimulus that was more likely to change in orientation. As expected, perceptual sensitivity increased for orientation changes in the cued stimulus location (Fig. 6*a* inset, red dot above black dot). ‘Attend-in’ trials were those in which the cued stimulus location was inside the aggregate receptive fields (RFs) of the recorded V4 neurons, whereas ‘attend-out’ trials were those in which the cued stimulus location was in the opposite visual hemifield.

**Figure 6:**
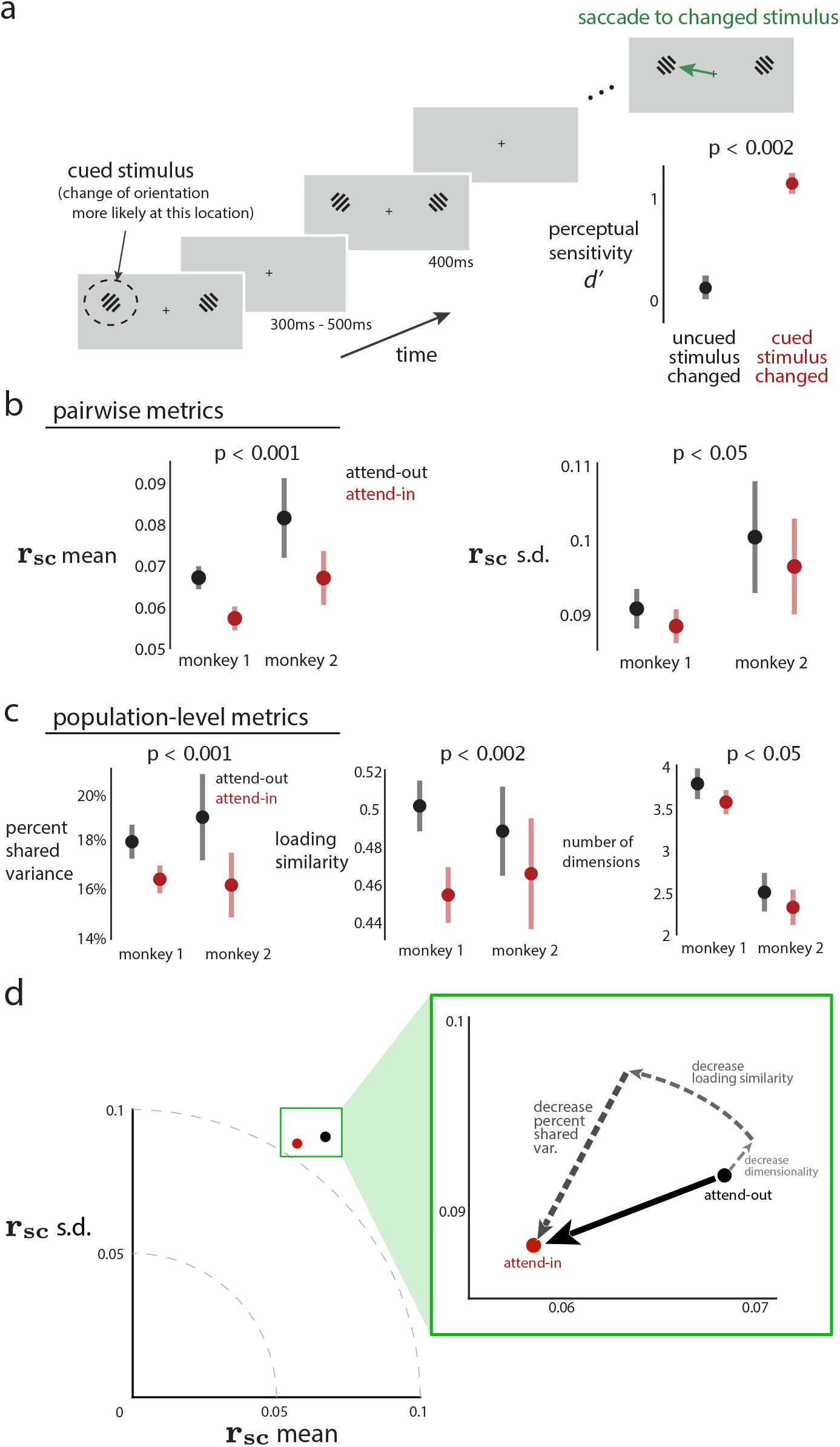
An observed decrease in *r*_sc_ mean of macaque V4 neurons during a spatial attention task corresponds to changes in multiple population metrics. **a**. Experimental task design. On each trial, monkeys maintained fixation while Gabor stimuli were presented for 400 ms (with 300-500 ms in between presentations). When one of the stimuli changed orientation, animals were required to saccade to the changed stimulus to obtain a reward. At the beginning of a block of trials, we performed an attentional manipulation by cuing animals to the location of the stimulus that was more likely to change for that block (dashed circle denotes the cued stimulus and was not presented on the screen). The cued location alternated between blocks. Animals were more likely to detect a change in stimulus at cued rather than uncued locations (inset in bottom right, *p* < 0.002 for both animals; data for monkey 1 is shown). During this task, we recorded activity from V4 neurons whose receptive fields (RFs) overlapped with one of the stimulus locations. **b**. *r*_sc_ mean (left panel) and *r*_sc_ s.d. (right panel) across recording sessions for two animals. Black denotes ‘attend-out’ trials (i.e., the cued location was outside the recorded V4 neurons’ RFs), and red denotes ‘attend-in’ trials (i.e., the cued location was inside the RFs). Data was pooled across both animals to compute *p*-values reported in titles for comparison of attend-out (black) and attend-in (red). For individual animals, *r*_sc_ mean was lower for attend-in than attend-out (*p* < 0.001 for each animal). *r*_sc_ s.d. was also lower for attend-in than attend-out (*p* < 0.05 for monkey 1, and *p* = 0.148 for monkey 2). **c**. Population metrics identified across recording sessions for two animals (same data as in **b**). Black denotes attend-in trials, red denotes attend-out trials. Data was again pooled across animals to compute *p*-values reported in titles for comparing attend-out and attend-in. %sv was lower for attend-in than attend-out (*p* < 0.001 for monkey 1 and *p* < 0.02 for monkey 2). Loading similarity was lower for attend-in than attend-out (*p* < 0.001 for monkey 1 and *p* = 0.162 for monkey 2). Dimensionality was lower for attend-in than attend-out (*p* = 0.113 for monkey 1 and *p* = 0.174 for monkey 2). In panels **a**-**c**, dots indicate means and error bars indicate 1 s.e.m., both computed across recording sessions. **d**. Summary of the real data results. Attention decreases both *r*_sc_ mean and *r*_sc_ s.d. (black dot to red dot). These decreases in pairwise metrics correspond to a combination of decreases in %sv, loading similarity, and dimensionality (dashed arrows).

For pairwise metrics, *r*_sc_ mean decreased when attention was directed into the RFs of the V4 neurons (Fig. 6*b*, left panel), consistent with previous studies (Cohen and Maunsell, 2009; Mitchell et al., 2009; Gregoriou et al., 2014; Snyder et al., 2016, 2018). We further found that *r*_sc_ s.d. was lower for attend-in trials than for attend-out trials, an effect not reported previously (Fig. 6*b*, right panel; also see Supplementary Fig. 1 for session-by-session pairwise metrics).

The decrease in both *r*_sc_ mean and *r*_sc_ s.d. could arise from several different types of distinct changes in population-wide covariability (Fig. 5). To compute the population metrics, we applied factor analysis (FA) separately to attend-out and attend-in trials (see Methods). FA is the most basic dimensionality reduction method that characterizes shared variance among neurons (Cunningham and Yu, 2014), and is consistent with how we created covariance matrices in Figures 3 and 4. We found three distinct changes in population metrics. First, neuronal activity during attend-in trials had lower %sv than during attend-out trials (Fig. 6*c*, left), consistent with previous interpretations that attention reduces the strength of shared modulations (Rabinowitz et al., 2015; Ecker et al., 2016; Huang et al., 2019; Ruff et al., 2019b). Second, we also found lower loading similarity for attend-in trials than attend-out trials for the dominant dimension (i.e., the dimension that explains the largest proportion of the shared variance; Fig. 6*c*, middle). This implies that, with attention, neurons in the population co-fluctuate in a more heterogeneous manner (i.e., more pairs of neurons co-fluctuate in opposition, and fewer pairs co-fluctuate together). Third, we found that dimensionality was slightly lower for attend-in than attend-out trials (Fig. 6*c*, right). Thus, on average, a smaller number of distinct shared signals were present when attention was directed into the neurons’ RFs. The small change in dimensionality is consistent with the relative strength of each dimension (i.e., eigenspectrum shape) being similar for attend-in and attend-out (Supplementary Fig. 2). Taken together, this collection of observations of both pairwise and population metrics leads to a more refined view of how attention affects population-wide covariability.

The pairwise (Fig. 6*b*) and population (Fig. 6*c*) metrics are computed based on the same recorded activity and each represents a different view of population activity. The central contribution of our work is to provide a framework by which to understand these two perspectives and five different metrics in a coherent manner. Using the relationships between pairwise and population metrics we have established in the *r*_sc_ mean versus *r*_sc_ s.d. space (Fig. 5), we can decompose the decrease in *r*_sc_ mean and s.d. into: 1) a small decrease in dimensionality (Fig. 6*d*, small dashed arrow), 2) a decrease in loading similarity (Fig. 6*d*, medium dashed arrow), and 3) a substantial decrease in %sv (Fig. 6*d*, large dashed arrow). Overall, this analysis demonstrates the insufficiency of any one measure of correlated variability, and the value in considering pairwise and population metrics together, with a bridge that allows one to navigate between the two.

## Discussion

Coordinated variability in the brain has long been linked to the neural computations underlying a diverse range of functions, including sensory encoding, decision making, attention, learning, and more. In this study, we sought to relate two major bodies of work investigating the coordinated activity among neurons: studies that measure spike count correlation between pairs of neurons (*r*_sc_) and studies that use dimensionality reduction to measure population-wide covariability. We considered three population metrics and established analytically and empirically that: 1) increasing loading similarity corresponds to increasing *r*_sc_ mean and decreasing *r*_sc_ s.d., 2) decreasing percent shared variance (%sv) corresponds to decreasing both *r*_sc_ mean and s.d., and 3) increasing dimensionality tends to decrease *r*_sc_ mean and s.d. Applying this understanding to recordings in macaque V4, we found that the previously-reported decrease in mean spike count correlation associated with attention stemmed from a decrease in %sv, a decrease in loading similarity, and decrease in dimensionality. This analysis revealed that attention involves multiple changes in how neurons interact that are not well captured by a single statistic alone. Overall, our work demonstrates that common ground exists between the literatures of spike count correlation and dimensionality reduction approaches, and builds the intuition and formalism to navigate between them.

Our work provides a cautionary tale for attempting to summarize population-wide covariability using one, or a small number of, statistics. For example, reporting only *r*_sc_ mean is incomplete because several distinct changes in population-wide covariability can correspond to the same change in *r*_sc_ mean. In a similar vein, reporting only dimensionality is incomplete because it does not indicate how strongly the neurons covary, nor their co-fluctuation patterns. For this reason, we recommend reporting several different pairwise and population metrics (e.g., the five used in this study along with the eigenspectrum of the shared covariance matrix), as long as they can be reliably measured from the data available. This not only allows for a deeper and more complete understanding of how neurons covary, but also it allows one to make tighter connections to previous literature that uses the same metrics. Future work may seek to revisit previous results of correlated neuronal variability that are based on a single statistic (e.g., *r*_sc_ mean), and reinterpret them within a framework that considers multiple perspectives and statistics of population-wide covariability, such as that presented here.

There are some situations where it is not feasible to reliably measure population statistics, such as recording from a small number of neurons in deep brain structures (Nevet et al., 2007; Liu et al., 2013). In such situations, the *r*_sc_ can be measured between pairs of neurons recorded in each session and then averaged across sessions to obtain the *r*_sc_ mean (Supplementary Fig. 3). Based on our findings, we recommend that studies which report *r*_sc_ mean also report *r*_sc_ s.d. because the latter provides additional information about population-wide covariability. For example, in the special case of one latent dimension (typically not known in advance for real data), measuring *r*_sc_ mean and *r*_sc_ s.d. allows one to estimate the loading similarity and %sv (cf. Fig. 3*e*-*f*). In general, even when there is more than one latent dimension in the population, *r*_sc_ s.d. provides value in situating the data in the *r*_sc_ mean versus *r*_sc_ s.d. plot (cf. Fig. 5). Changes in *r*_sc_ mean and s.d. can then inform changes in population metrics based on the relationships established in this work (cf. Fig. 6*d*).

We considered three population metrics — dimensionality, percent shared variance (%sv), and loading similarity — that summarize the structure of population-wide covariability and are rooted in well-established concepts in existing literature. First, dimensionality has been used to describe how neurons covary across conditions (i.e., an analysis of trial-averaged firing rates; Churchland et al., 2012; Rigotti et al., 2013; Mante et al., 2013; Cowley et al., 2016; Kobak et al., 2016; Sohn et al., 2019), as well as how neurons covary from trial to trial (Yu et al., 2009; Santhanam et al., 2009; Sadtler et al., 2014; Rabinowitz et al., 2015; Mazzucato et al., 2016; Williamson et al., 2016; Bittner et al., 2017; Athalye et al., 2017; Williams et al., 2018; Stringer et al., 2019a; Recanatesi et al., 2019). We focused on the latter in our study to connect with the *r*_sc_ literature, which also seeks to understand the shared trial-to-trial variability between neurons. To focus on the shared variability among neurons, we used factor analysis (FA) to measure dimensionality. Another commonly-used dimensionality reduction method, principal components analysis (PCA), although appropriate for studying trial-averaged activity, does not distinguish between variability that is shared among neurons and variability that is independent to each neuron. Second, investigating the loading similarity has provided insight about whether shared variability among neurons arises from a shared global factor which drives neurons to increase and decrease their activity together (Ecker et al., 2014; Okun et al., 2015; Lin et al., 2015; Rabinowitz et al., 2015; Williamson et al., 2016; Huang et al., 2019) or whether the co-fluctuations involve a more intricate pattern across the neuronal population (Snyder et al., 2018; Insanally et al., 2019; Cowley et al., 2020). Understanding how loading similarity and these patterns of shared variability interact with (e.g., align with or are orthogonal to) patterns of stimulus encoding and downstream readouts will be important to understand how the brain perceives and computes (Averbeck et al., 2006; Moreno-Bote et al., 2014; Kohn et al., 2016; Ni et al., 2018; Ruff and Cohen, 2019a; Cowley et al., 2020; Rumyantsev et al., 2020; Bartolo et al., 2020). Third, we have previously reported %sv for area V1 (Williamson et al., 2016), area M1 (Hennig et al., 2018), and network models (Williamson et al., 2016; Bittner et al., 2017). Conceptually, %sv and *r*_sc_ mean are both designed to capture the strength of shared variability in a population of neurons. Thus, we might initially think that there should be a one-to-one correspondence between the two quantities. Indeed, if the population activity is described by one co-fluctuation pattern with a high loading similarity, there is a direct relationship between %sv and *r*_sc_ mean (Fig. 3*f*). However, in general, %sv and *r*_sc_ mean do not have a one-to-one correspondence between them (Fig. 3*f*, moderate or low loading similarity).

Although pairwise correlation and dimensionality reduction have most commonly been computed based on spike counts, several studies have also computed these metrics on neuronal activity recorded using other modalities, such as calcium imaging (Harvey et al., 2012; Ahrens et al., 2012; Dechery and MacLean, 2018; Stringer et al., 2019a; Rumyantsev et al., 2020). The relationships that we established here between pairwise and population metrics are properties of covariance matrices in general and do not rely on or assume recordings of neuronal spikes. Thus, the intuition built here can be applied to other recording modalities.

Our work here focused on studying interactions within a single population of neurons. Technological advances are enabling recordings from multiple distinct populations simultaneously, including neurons in different brain areas, neurons in different cortical layers, or different neuron types (e.g., Ahrens et al., 2013; Jiang et al., 2015; Jun et al., 2017). Studies are dissecting the interactions between these distinct populations using pairwise correlation (Smith et al., 2012; Pooresmaeili et al., 2014; Oemisch et al., 2015; Zandvakili and Kohn, 2015; Ruff and Cohen, 2016a; Snyder et al., 2016) and dimensionality reduction (Semedo et al., 2014; Buesing et al., 2014; Bittner et al., 2017; Perich et al., 2018; Semedo et al., 2019; Ames and Churchland, 2019; Ruff and Cohen, 2019a; Veuthey et al., 2020; Cowley et al., 2020). As we have shown here for a single population of neurons, considering a range of metrics from both the pairwise correlation and dimensionality reduction perspectives, and understanding how they relate to one another, will provide rich descriptions of how different neuronal populations interact.

## Data and code availability

Relevant data and analysis code used to generate the results are available upon request from the authors.

## Acknowledgements

The authors would like to thank Samantha Schmitt for help with data collection, and João Semedo for helpful discussions. BRC was supported by CV Starr Foundation Fellowship. ACS was supported by NIH grant K99EY025768. MAS and BMY were supported by NIH CRCNS R01 MH118929, NIH R01 EB026953, and NSF NCS BCS 1954107/1734916. MAS was supported by NIH R01 EY022928, NIH R01 EY029250, NIH P30 EY008098. BMY was supported by Simons Foundation 364994 and 543065, NIH R01 HD071686, NIH CRCNS R01 NS105318, and NSF NCS BCS1533672.

## Author contributions

Mathematical derivations: AU, RM; simulations: AU, RM, BRC; data collection: ACS; data analysis: AU, BRC, writing: AU, RM, BRC; reviewing and editing: all authors. AU, RM, BRC contributed equally to this work (first authors). MAS, BMY contributed equally to this work (senior authors).

## Competing interests

The authors declare no competing interests.

## Methods

### Spike count covariance matrix

Both pairwise metrics and population metrics are computed directly from the spike count covariance matrix Σ of size *n* × *n* for a population of *n* neurons. Each entry in Σ is the covariance between the activity of neuron *i* and neuron *j*:

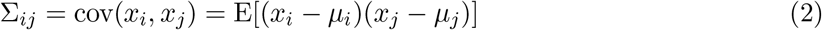

where *x*_*i*_ and *x*_*j*_ represent the activity of neurons *i* and *j*, respectively, and *μ*_*i*_ and *μ*_*j*_ represent the mean activity of neurons *i* and *j*, respectively. The variance of the *i*th neuron is equal to Σ_*ii*_.

### Pairwise metrics

We computed the spike count correlation (*r*_sc_) between neurons *i* and *j* directly from the spike count covariance matrix:

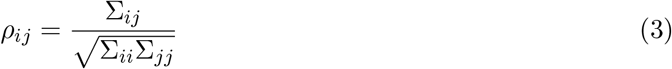

We then summarized the distribution of *r*_sc_ values across all pairs of neurons in the population with two pairwise metrics: the *r*_sc_ mean and *r*_sc_ standard deviation (s.d.).

### Population metrics

The metrics we use for characterizing population-wide covariability are based on factor analysis (FA; Santhanam et al., 2009; Yu et al., 2009; Churchland et al., 2010; Harvey et al., 2012; Williamson et al., 2016; Bittner et al., 2017; Athalye et al., 2017; Huang et al., 2019), a dimensionality reduction method. We chose FA because it is the most basic dimensionality reduction method that explicitly separates variance that is shared among neurons from variance that is independent to each neuron. This allows us to relate the population metrics provided by FA to spike count correlation, which is designed to measure shared variability between pairs of neurons. One might consider using principal component analysis (PCA), but it does not distinguish shared variance from independent variance. Thus, FA is more appropriate than PCA for studying the shared variability among a population of neurons.

#### Decomposing the spike count covariance matrix

FA decomposes the spike count covariance matrix Σ into a low-rank shared covariance matrix, which captures the variability shared among neurons in the population, and an independent variance matrix, which captures the portion of variance of each neuron unexplained by the other neurons (Fig. 7*a*):

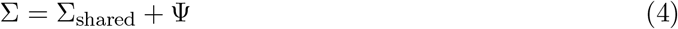

where Σ_shared_ ∈ ℝ^*n×n*^ is the shared covariance matrix for *n* neurons, and ψ ∈ ℝ^*n×n*^ is a diagonal matrix containing the independent variance of each neuron. The low-rank shared covariance matrix can be expressed using the eigendecomposition as (Fig. 7*a*):

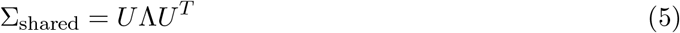

where *U* ∈ ℝ^*n×d*^ and Λ ∈ ℝ^*d×d*^, with *d < n*. The rank (i.e., dimensionality) of the shared covariance matrix, *d*, indicates the number of latent variables. Each column of *U* is an eigenvector and represents a co-fluctuation pattern containing the loading weights of each neuron (i.e., how much each neuron contributes to that dimension). The matrix Λ is a diagonal matrix where each diagonal element is an eigenvalue and represents the amount of variance along the corresponding co-fluctuation pattern (e.g., in Fig. 2 panel *a* has larger eigenvalue than panel *c*).

**Figure 7:**
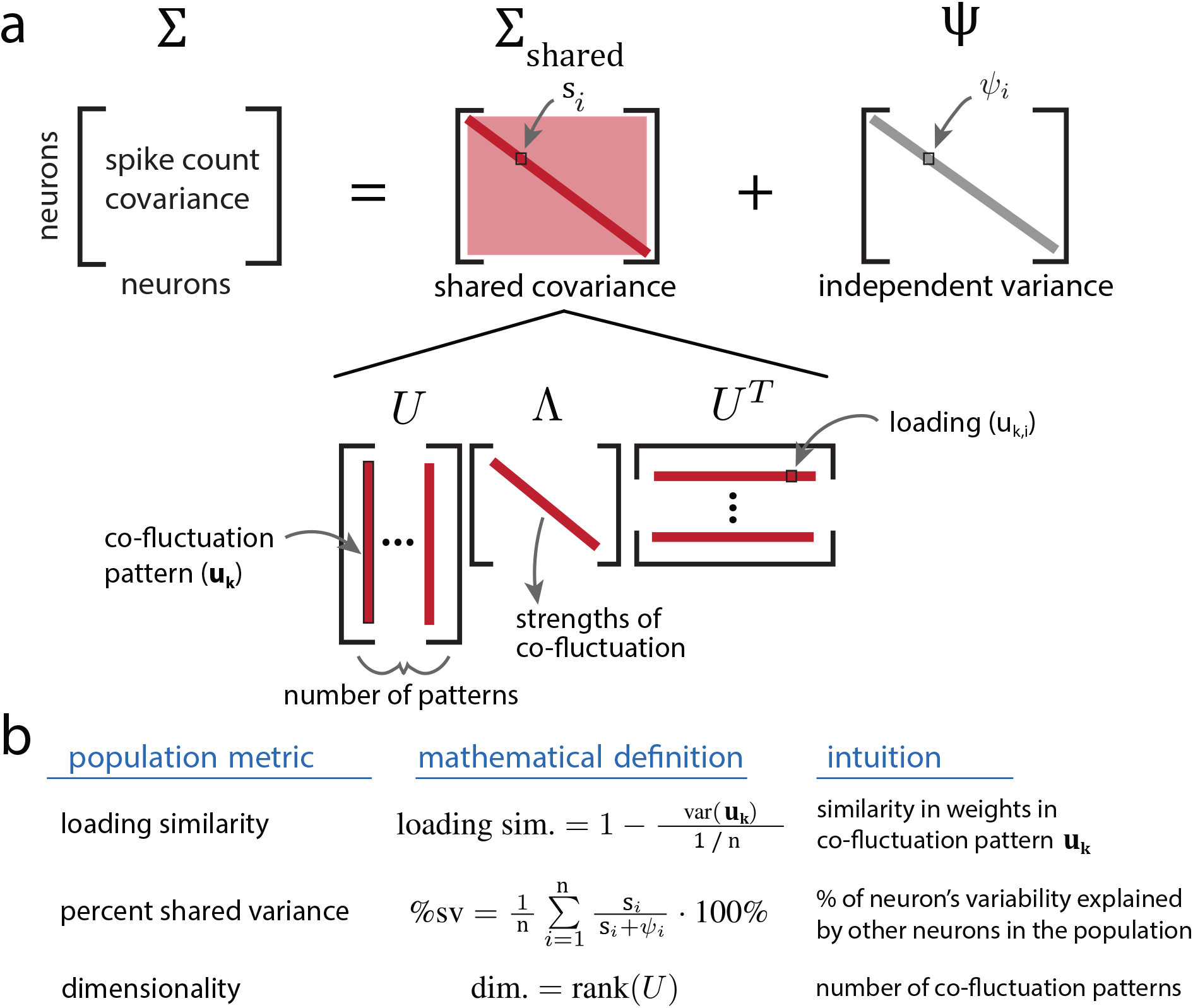
Decomposition of the spike count covariance matrix and defining population metrics. **a**. We use factor analysis to decompose the spike count covariance matrix Σ into the sum of a low-rank shared covariance matrix Σ_shared_ and a diagonal independent variance matrix ψ. The *i*th diagonal entry of Σ_shared_ (*s*_*i*_) corresponds to the spike count variance that neuron *i* shares with other neurons in the population (i.e., shared variance), while the *i*th diagonal entry of ψ_*i*_ corresponds to spike count variance of neuron *i* that cannot be explained by the other neurons (i.e., independent to neuron *i*). We can further decompose Σ_shared_ via an eigendecomposition to extract the co-fluctuation patterns (i.e., the eigenvectors) and the strength of each latent co-fluctuation (i.e., the eigenvalues). **b**. The population metrics used in this study are loading similarity, percent shared variance (%sv), and dimensionality.

Based on this matrix decomposition, we defined the three metrics that describe the population-wide covariability:

- **Loading similarity**: the similarity of loading weights across neurons for a given co-fluctuation pattern. Scalar value between 0 (the weights are maximally dissimilar, defined precisely below) and 1 (all weights are the same).
- **Percent shared variance (%sv)**: the percentage of each neuron’s variance that is explained by other neurons in the population. Percentage between 0% and 100%.
- **Dimensionality**: the number of dimensions (i.e., co-fluctuation patterns). Integer value.

We give the precise definitions of these population metrics below and in Fig. 7*b*.

#### Loading similarity

We sought to define loading similarity such that, for a given co-fluctuation pattern, if the weights for all neurons are the same, we would measure a loading similarity of 1. When the weights are as different as possible, we would measure a loading similarity of 0. We define the loading similarity based on the variance across the *n* weights (for *n* neurons) in a co-fluctuation pattern **u**_**k**_. The smallest possible variance is 0; the largest possible variance, for a unit vector **u**_**k**_, is 1*/n* (Supplementary Math Note F). Thus, we define loading similarity for a co-fluctuation pattern **u**_**k**_ ∈ ℝ^*n*^ as:

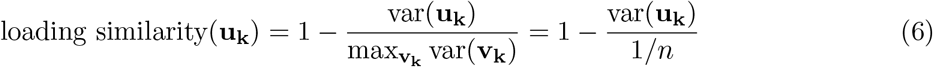

where the loading similarity is computed on unit vectors (i.e., **u**_**k**_ has a norm of 1). The notation var(**u**_**k**_) denotes that the variance is being taken across the *n* elements of the vector **u**_**k**_. The denominator of equation (6) acts as a normalizing factor, bounding the loading similarity value between 0 and 1.

The loading similarity distinguishes between a co-fluctuation pattern along which all neurons in the population have the same weight in which case they change their activity up and down together (Fig. 2*a*; loading similarity of 1), from one in which weights are different and some neurons increase their activity when others decrease their activity (Fig. 2*b*; loading similarity of 0). The loading weights we use here are closely related to ‘population coupling’ (Okun et al., 2015) and ‘modulator weights’ (Rabinowitz et al., 2015). For some types of shared fluctuations, these weights are similar across neurons in a population (i.e., high loading similarity; Okun et al., 2015; Rabinowitz et al., 2015; Huang et al., 2019). For other types of shared fluctuations, the weights vary substantially across neurons in the population (i.e., low loading similarity; Snyder et al., 2018; Cowley et al., 2020).

We show in Supplementary Math Note E why, if one dimension has high loading similarity, the other dimensions must have low loading similarity. The reason is that co-fluctuation patterns are defined to be mutually orthogonal. If one co-fluctuation pattern has all weights close to the same value (i.e., high loading similarity), then all other co-fluctuation patterns must have substantial diversity in their weights (i.e., low loading similarity) to satisfy orthogonality.

#### Percent shared variance

The percent shared variance (%sv) measures the percentage of each neuron’s spike count variance that is explained by other neurons in the population (Williamson et al., 2016; Bittner et al., 2017; Hennig et al., 2018). Equivalently, we can think of %sv in terms of latent co-fluctuations. Because latent co-fluctuations capture the shared variability among neurons, the %sv measures how much of each neuron’s variance is explained by the latent co-fluctuations. The activity of neurons may be tightly linked to the latent co-fluctuation (e.g., Fig. 2*a*), in which case a large percentage of each neuron’s variance is shared with other neurons, or may only be loosely linked to the latent co-fluctuation (e.g., Fig. 2*c*), in which case a small percentage of each neuron’s variance is shared with other neurons. Mathematically, we define the %sv for a neuron *i*:

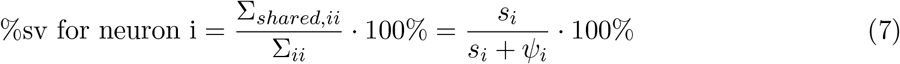

where *s*_*i*_ is the *i*^*th*^ entry along the diagonal of the shared covariance matrix (Fig. 7*a*, Σ_shared_), and *ψ*_*i*_ is the *i*^*th*^ entry along the diagonal of the independent covariance matrix (Fig. 7*a*, ψ). A %sv of 0% indicates that the neuron does not covary with (i.e., is independent of) other neurons in the population, whereas a %sv of 100% indicates that the neuron’s activity can be entirely accounted for by the activity of other neurons in the population. To compute %sv for an entire population of neurons, we averaged the %sv of the individual neurons. All %sv values reported in this study are the %sv for the neuronal population.

#### Dimensionality

Dimensionality refers to the number of latent co-fluctuations needed to describe population-wide covariability. For example, the population-wide covariability can be described by one latent co-fluctuation (Fig. 2*a*) or by several latent co-fluctuations (Fig. 2*d*). In the population activity space, dimensionality corresponds to the number of axes along which the population activity varies (see Fig. 2*d*, bottom inset). Mathematically, the dimensionality is the rank of the shared covariance matrix (i.e., the number of columns in *U*, Fig. 7*a*).

### Creating the spike count covariance matrices with specified population metrics

To relate pairwise and population metrics, we created spike count covariance matrices of the form in equation (4) with specified population metrics. Importantly, we did not simulate spike counts, nor fit a factor analysis model to simulated data. Rather, we created covariance matrices using (4) and computed pairwise correlations directly from the entries of the covariance matrix, as shown in (3). Across simulations (Figs. 3 and 4), we simulated with *n* = 30 neurons and set independent variances (i.e., diagonal elements of ψ in equation (4)) to 1.

#### Specifying co-fluctuation patterns to obtain different loading similarities

Each co-fluctuation pattern **u**_**k**_ is a vector with *n* = 30 entries (one entry per neuron). We generated a single co-fluctuation pattern by randomly drawing 30 independent samples from a Gaussian distribution with a mean of 2.5. We choose a nonzero mean so that we could obtain co-fluctuation patterns with loading similarities close to 1 when drawing from the Gaussian distribution (i.e., a mean of 0 would have resulted in almost all co-fluctuation patterns having a loading similarity close to 0). To get a range of loading similarities between 0 and 1, we used different standard deviations for the Gaussian. For a small standard deviation value, all entries in the co-fluctuation pattern are close to 2.5, resulting in a high loading similarity. For larger standard deviations, some loading weights are positive and some negative, with large variability in their values, resulting in co-fluctuation patterns with low loading similarity. We increased the Gaussian standard deviation from 0.1 to 5.5 with increments of size 0.1. For each increment, we generated 50 patterns and normalized them to have unit norm. In total, we created a set of 2,750 random patterns.

The following procedure describes the construction of shared covariance matrices with one co-fluctuation pattern. We chose a single pattern **u**_**1**_ ∈ ℝ^30×1^ (i.e., *U* has only 1 column) from the set of 2,750. We constructed the shared covariance matrix by computing *U* Λ*U*^*T*^, where Λ was chosen to achieve a desired percent shared variance (see below). The covariance matrix was then computed according to equation (4). We created a covariance matrix, yielding a spread of loading similarities between 0 and 1 (Fig. 3*e*-*f*). In the next section, we describe the procedure for creating a covariance matrix with more dimensions.

#### Specifying the percent shared variance

To achieve a given %sv, either the independent variance or the amount of shared variability (i.e., the eigenvalues) of each dimension can be adjusted. In the main text, we set the independent variance of each neuron to ψ_*i*_ = 1, and changed the total amount of shared variability by multiplying each eigenvalue (each diagonal element in Λ from equation (5)) by the same constant value, *a*. To obtain a specified %sv, we identified *a* by searching through a large set of possible values (from 10^*−*4^ to 10^3^ with step size 10^*−*3^). We allowed for a tolerance of *ϵ* = 10^*−*3^ between the desired %sv and the %sv that was achieved after scaling the eigenvalues by *a*. In Supplementary Figure 4, we allowed the independent variances to be different across neurons, and the results were qualitatively similar to that shown in the main text.

#### Increasing dimensionality

To assess how changing dimensionality affects pairwise metrics, we created covariance matrices whose shared covariance matrix comprised more than 1 dimension. To create a shared covariance matrix with *d* dimensions, we randomly chose *d* patterns from the set of 2750 we had generated above (see ‘Specifying co-fluctuation patterns to obtain different loading similarities’). We then orthogonalized the chosen patterns using the Gram-Schmidt process to obtain *d* orthonormal (i.e., orthogonal and unit length) co-fluctuation patterns *U* ∈ ℝ^30×*d*^. We formed the shared covariance matrix using *U* Λ*U*^*T*^, where Λ ∈ ℝ^*d×d*^ is a diagonal matrix containing the eigenvalues (i.e., the strength of each dimension; see ‘Specifying the relative strengths of each dimension’ below). We repeated this procedure to produce 3,000 sets of *d* orthonormal patterns (i.e., 3,000 different *U* matrices), each of which was used to create a shared covariance matrix. The spike count covariance was computed according to equation (4).

#### Specifying the relative strengths of each dimension

In simulating shared covariance matrices with more than one dimension, we chose the relative strength of each dimension by specifying the eigenspectrum (diagonal elements of Λ in equation (5)). We worked with three sets of eigenspectra. First, a flat eigenspectrum had eigenvalues that were all equal (Fig. 3*g*). Second, for two dimensions, we varied the ratio of the two eigenvalues between 95:5, 80:20, 50:50, 20:80, and 5:95 (Fig. 4). Third, we considered an eigenspectrum in which each subsequent eigenvalue falls off according to an exponential function (Supplementary Fig. 5). Only the relative (and not the absolute) eigenvalues (i.e., the shape of the eigenspectrum) affect the results, because the eigenspectrum was subsequently scaled to achieve a desired %sv (see ‘Specifying the values of percent shared variance’).

### Analysis of V4 neuronal recordings from a spatial attention task

#### Electrophysiological recordings

We analyzed data from a visual spatial attention task reported in a previous study (Snyder et al., 2018). Briefly, we implanted a 96-electrode “Utah” array (Blackrock Microsystems; Salt Lake City, UT) into visual cortical area V4 of an adult male rhesus macaque monkey (data from two monkeys were analyzed; in our study, monkey 1 corresponds to “monkey P” and monkey 2 corresponds to “monkey W” from Snyder et al. (2018)). After recording electrode voltages (Ripple Neuro.; Salt Lake City, UT), we used custom software to perform off-line spike sorting (Kelly et al., 2007, freely available at https://github.com/smithlabvision/spikesort). This yielded 93.2 ± 8.9 and 61.9 ± 27.4 candidate units per session for monkey 1 and 2, respectively.

To further ensure the isolation quality of recorded units, we removed units from our analyses according to the following criteria. First, we removed units with a signal-to-noise ratio of the spike waveform less than 2.0 (Kelly et al., 2007). Second, we removed units with overall mean firing rates less than 1 Hz, as estimates of *r*_sc_ for these units tends to be poor (Cohen and Kohn, 2011). Third, we removed units that had large and sudden changes in activity due to unstable recording conditions. For this criterion, we divided the recording session into ten equally-sized blocks and for each unit computed the difference in average firing rate between adjacent blocks. We excluded units with a change in average firing rate greater than 60% of the maximum firing rate (where the maximum is taken across the ten equally-sized blocks). Fourth, we removed an electrode from each pair of electrodes that were likely electrically-coupled. We identified the coupled electrodes by computing the fraction of threshold crossings that occurred within 100 *μ*s of each other for each pair of electrodes. We then removed the fewest number of electrodes to ensure this fraction was less than 0.2 (i.e., pairs with an unusually high number of coincident spikes) for all pairs of electrodes. Fifth, we removed units that did not sufficiently respond to the visual stimuli used in the experiment. Evoked spike counts (i.e., a neuron’s response after stimulus presentation) were taken between 50 ms to 250 ms after stimulus onset, and spontaneous spike counts (i.e., a neuron’s response during a blank screen) were taken in a 200 ms window that ended 50 ms before stimulus onset. For each unit, we computed a sensitivity measure *d′* between evoked and spontaneous activity:

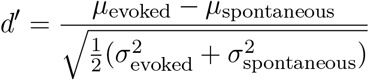

for mean spike counts *μ*_evoked_ and *μ*_spontaneous_ and spike count variances 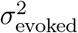 and 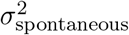. We removed units with *d′* < 0.5 from analyses, as these units had spontaneous and evoked responses that were difficult to distinguish.

After applying these five criteria, 44.5 ± 11.3 and 18.8 ± 6.7 units per session (mean ± s.d. over sessions) remained for monkeys 1 and 2, respectively. Although these remaining units likely contained both single-unit and multi-unit activity, we refer to each unit as a neuron for simplicity.

#### Visual stimulus change-detection task

Animals were trained to perform a change-detection task with a spatial attention cue to the location of the visual stimulus that was more likely to change (Snyder et al., 2018). In the visual change-detection task (Fig. 6*a*), animals fixated a central dot while Gabor stimuli were presented in two locations on a computer screen. One location was chosen to be within the aggregate receptive fields (RFs) of the recorded V4 neurons (mapped prior to running the experiment), and the other location was placed at the mirror symmetric location in the opposite hemifield. Animals maintained fixation while a sequence of Gabor stimuli were presented. Each drifting Gabor stimulus (oriented at either 45° or 135°) was presented for 400 ms, followed by a blank screen presented for a random interval (between 300 and 500 ms). The sequence continued, with a fixed probability for each presentation, until one of the two stimuli changed orientation when presented (i.e., the ‘target’). Upon target presentation, animals were required to make a saccade to the target to earn a juice reward. We manipulated spatial attention in the experiment by cueing the more probable target location in blocks. At the beginning of each block, the cue was denoted by presenting only one Gabor stimulus at the more probable target location (90% likely), and requiring animals to detect orientation changes at this location for 5 trials. Consistent with the results of previous studies, we found that animals had greater perceptual sensitivity for orientation changes at the cued (i.e., attended) location than the uncued location (Fig. 6*a*, inset in the bottom right) and shorter reaction times (Snyder et al., 2018).

#### Data processing and computing spike counts

We first separated the trials into two groups: (1) “attend in” trials, for which the cued stimulus was inside the recorded neurons’ RFs and (2) “attend out” trials, for which the cued stimulus was outside the RFs. Since the initial orientation of the stimulus at the cued location could be one of two values (i.e., 45° or 135°), we further divided trials, resulting in a total of 4 groups of trials per session (attend in & 45°, attend out & 45°, attend in & 135°, attend out & 135°). Each combination of cued location and stimulus orientation was treated as an independent sample. The same neurons were used for each of the 4 groups within each session, ensuring a fair comparison between the attend-in and attend-out conditions.

We analyzed all stimulus presentations for which the target stimulus did not change. For each stimulus presentation, we took spike counts in a 200 ms window starting 150 ms after stimulus onset. For each of the 4 groups, we formed a spike count matrix *X* ∈ ℝ^*n×t*^, containing the spike counts of the *n* recorded neurons for the *t* trials belonging to that group. These spike count matrices were then used to compute both the pairwise and population metrics (described below). For all analyses (Fig. 6), we excluded recording sessions with fewer than 10 neurons. Additionally, because population metrics depend on the number of trials (Williamson et al., 2016), for each session we equalized the number of trials across the 4 groups by randomly subsampling from groups with larger numbers of trials.

#### Computing pairwise metrics for V4 spike counts

We computed pairwise metrics on each combination of attention state (‘attend in’ and ‘attend out’) and stimulus orientation. We computed the correlation matrix for *X* as described above in ‘Pairwise metrics’ and then computed *r*_sc_ mean and *r*_sc_ s.d. For each attention state, we averaged the *r*_sc_ mean and *r*_sc_ s.d. over sessions and different stimulus orientations.

#### Computing population metrics for V4 spike counts

We fit the parameters of a factor analysis model (see Fig. 7*a*) to each spike count matrix *X* (as described above) using the expectation-maximization (EM) algorithm (Dempster et al., 1977). For each session, this was performed separately for each attention state and stimulus orientation. Using the FA parameters, we then computed the three population metrics (Fig. 7*b*). For dimensionality, we first found the number of dimensions *d* that maximized the cross-validated data likelihood. We fit an FA model with *d* dimensions, and then found the number of dimensions required to explain 95% of the shared variance, termed *d*_*shared*_ (Williamson et al., 2016). We report *d*_*shared*_ because it tends to be a more reliable estimate of dimensionality than the number of dimensions that maximizes the cross-validated data likelihood. We computed %sv as described by equation (7). We report the loading similarity as defined in equation (6) for the co-fluctuation pattern that explained the most shared variability (i.e., the eigenvector with the largest eigenvalue), since it contributes most to describing the population-wide covariability. For ‘attend in’ and ‘attend out’ conditions, we averaged the population metrics across sessions and stimulus orientations.

### Statistics

We employed paired permutations tests for all statistical comparisons of pairwise metrics and population metrics between ‘attend-in’ and ‘attend-out’ conditions (Fig. 6*b*-*c*). First, for a given, metric we computed the average difference between attend-in and attend-out. Then, we computed a null distribution by randomly permuting attend-in and attend-out labels and recomputing the average difference in the permuted data. We ran 10,000 permutations to obtain a null distribution of 10,000 samples. We computed *p*-values as the proportion of samples in the null distribution that were more extreme than the average difference in the data, corresponding to *p* < 0.0001 as the highest attainable level of significance in our statistical analyses.

#### Supplementary Math Notes

##### A Relationship between correlation, loading similarity, and %sv (one latent dimension)

We establish here the mathematical relationship between *r*_sc_, loading similarity, and %sv. This will provide the formalism for understanding why decreasing %sv decreases both *r*_sc_ mean and s.d. (Fig. 3*f*), that a high loading similarity corresponds to large *r*_sc_ mean and low *r*_sc_ s.d. (Fig. 3*e*), and that a low loading similarity corresponds to small *r*_sc_ mean and large *r*_sc_ s.d. (Fig. 3*e*).

Let *n* be the number of neurons, and let **w** be the co-fluctuation pattern (i.e., loading vector [*w*_1_, *w*_2_, …, *w*_*n*_]^*T*^ ∈ ℝ^*n*×1^), *λ* ∈ ℝ_+_ be the strength of the co-fluctuation pattern (i.e., eigenvalue of the shared covariance matrix), and ψ ℝ^*n×n*^ be a diagonal matrix specifying the independent variance of each neuron (*ψ*_1_, *ψ*_2_, …, *ψ*_*n*_). Then the covariance matrix of the population activity is (see Methods and Fig. 7):

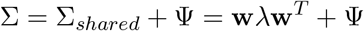

From this, we observe that Σ_*ij*_ = Σ_*shared,ij*_ = *λw*_*i*_*w*_*j*_ on the off-diagonal entries (i.e., if *i* ≠ *j*). Along the diagonals, 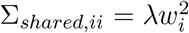 and 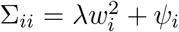. The correlation (i.e., *r*_sc_ if Σ is a spike count covariance matrix) between neurons *i* and *j* can be written as:

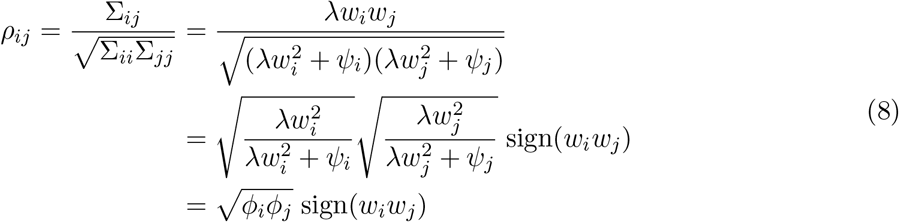

where *ϕ*_*i*_ and *ϕ*_*j*_ represent the %sv (as proportions) for neurons *i* and *j*, respectively, and sign(*w*_*i*_*w*_*j*_) = +1 if *w*_*i*_*w*_*j*_ > 0 or −1 if *w*_*i*_*w*_*j*_ < 0. The last line follows from the fact that %sv is defined in equation (7) as:

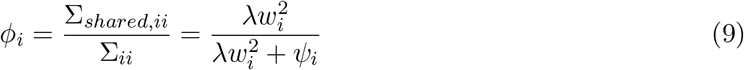

Equations (8) and (9) provide a basis for understanding the relationships between *r*_sc_, %sv, and loading similarity. The *r*_sc_ mean and s.d. are computed across all pairs of neurons *ρ*_*ij*_, for *i < j*.

For establishing a relationship between pairwise metrics and %sv, consider decreasing the overall %sv of the population, while keeping the loadings *w*_*i*_ fixed. This correspond o decreasing *λ* in equation (9), which implies *ϕ*_*i*_ for each neuron decreases, and thus the product 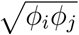 decreases for all pairs. The magnitude of each *ρ*_*ij*_ decreases (i.e., each *ρ*_*ij*_ moves closer to 0). As such, decreasing %sv of the population decreases the distance of a point from the origin in the *r*_sc_ mean versus *r*_sc_ s.d. plot, all else being equal (Fig. 3*f*).

For establishing a relationship between pairwise metrics and loading similarity, consider two extreme cases: 1) when loading similarity is 1 (as high as possible) 2) when it is 0 (as low as possible). We first assume that each neuron has the same independent variance *ψ*_*i*_ for simplicity, as we did in Figure 3. A loading similarity of 1 corresponds to each 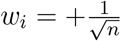 or each 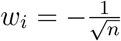. In either case, sign(*w*_*i*_*w*_*j*_) is always +1. Furthermore, *ϕ*_*i*_ is the same for every neuron and 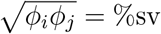 (i.e., the %sv of the population, expressed as a proportion) for every pair of neurons. Thus, all *ρ*_*ij*_ = %sv for all pairs of neurons *i* and *j*. In this case, *r*_sc_ mean = %sv and *r*_sc_ s.d. = 0. If the independent variances *ψ*_*i*_ are different across neurons, we can still √get each sign(*w*_*i*_*w*_*j*_) = +1 and each *ϕ*_*i*_ to be the same by setting each 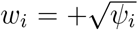 *ψ*_*i*_ or each 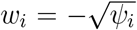. This would also result in *ρ*_*ij*_ = %sv for all pairs of neurons *i* and *j*, and thus *r*_sc_ mean = %sv and *r*_sc_ s.d. = 0. In this case, the loading similarity is still high (all *w*_*i*_ are the same sign; we can show that load. sim.*>* 0.5), but not equal to 1.

Now, consider a scenario in which half the loadings are 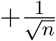 and the other half are 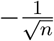 (and assume again that *ψ*_*i*_ are the same for every neuron). This is one way to obtain a loading similarity of 0. In this case, *ϕ*_*i*_ are still the same for every neuron, so 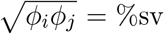 for all pairs. However, sign(*w*_*i*_*w*_*j*_) = −1 for 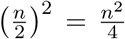 pairs, and sign(*w*_*i*_*w*_*j*_) = +1 for 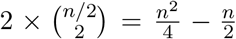 pairs. We can show that *r*_sc_ 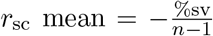 and, by using equation (10) from Supplementary Math Note B below, *r*_sc_ 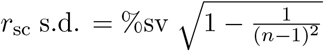. Thus, for a large number of neurons *n*, this case (where loading similarity=0) corresponds to small negative *r*_sc_ mean (close to 0), and large *r*_sc_ s.d. (close to the %sv). As an example, for 30 neurons and %sv=50%, this corresponds to *r*_sc_ mean = −0.0172 and *r*_sc_ s.d. = 0.4997.

With this analysis, we have established that for one latent dimension:

- Decreasing %sv decreases the magnitudes of correlations (i.e., each *ρ*_*ij*_ closer to 0). *r*_sc_ mean and s.d. both decrease (as seen empirically in Fig. 3*f*).
- Starting from a loading similarity near 1, a decrease in loading similarity involves flips in the signs of some correlations (i.e., some *ρ*_*ij*_ become −*ρ*_*ij*_). *r*_sc_ mean decreases but *r*_sc_ s.d. increases (as seen empirically in Fig. 3*f*).
- Both *r*_sc_ mean and %sv measure shared variance among neurons, but they are not always equal. Equations (8) shows that the two quantities are equal if all sign(*w*_*i*_*w*_*j*_) are the same (i.e., when loading similarity is high). However, in general *r*_sc_ mean and shared variance (%sv) are not the same—e.g., when loading similarity is low, or when there are multiple dimensions (Supplementary Math Note C).

In this section, we consider the extremes of loading similarity. In the next section, we analyze how gradual changes in loading similarity affect *r*_sc_ mean and s.d. for a fixed %sv.

##### B Circular arc in *r*_*sc*_ mean versus *r*_*sc*_ s.d. plot for one latent dimension and fixed %sv

We establish here mathematically that gradually varying the loading similarity for one latent dimension and fixed %sv results in an arc-like relationship between *r*_sc_ mean and *r*_sc_ s.d., and that the radius of the arc is approximately equal to the %sv (Fig. 3*e*-*f*).

We use the same notation as in Supplementary Math Note A. Let *E*[.] and *V ar*(.) denote the mean and variance across all neurons or all pairs of neurons, depending on context. In particular, we are interested in *E*[*ρ*] = *r*_sc_ mean, 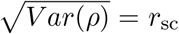 s.d., where the expectation and variance are computed across *ρ*_*ij*_ for all pairs of neurons in a given population (i.e., the upper triangle of the correlation matrix, *ρ*_*ij*_ for *i > j*).

Let *c* be the distance of a point (corresponding to one instance of the population activity covariance matrix) from the origin in the *r*_sc_ mean versus *r*_sc_ s.d. plot (i.e., 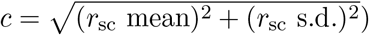). We want to know whether *c* is the same for all population activity covariance matrices with one latent dimension and fixed %sv. This would correspond to point being equidistant from the origin, and thus a circular arc. We can write *c* as:

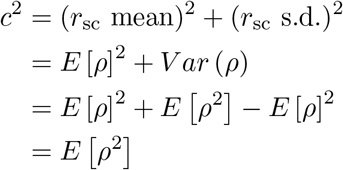

Thus, the squared distance (i.e., squared radius) is equal to *E* [*ρ*]^2^ an of 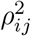 across all pairs in the population. Let *m* be the number of pairs (i.e., 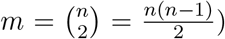). Now, using equations (8) and (9) derived in Supplementary Math Note A:

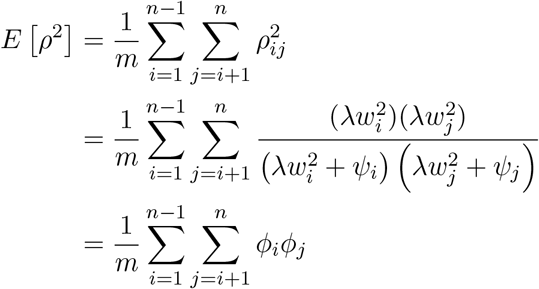

where *ϕ*_*i*_ and *ϕ*_*j*_ are the %sv of neurons *i* and *j* (expressed as proportions), as defined in Supplementary Math Note A. We can show that 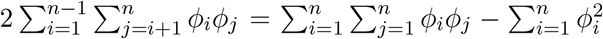. Intuitively, if we have a symmetric matrix ϕ with entries ϕ(*i, j*) = *ϕ*_*i*_*ϕ*_*j*_, and we want to find the sum of the off-diagonal elements 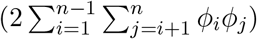, then we can take the sum of all elements and subtract the diagonal elements 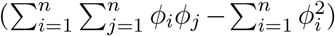. Using this equivalence, it follows:

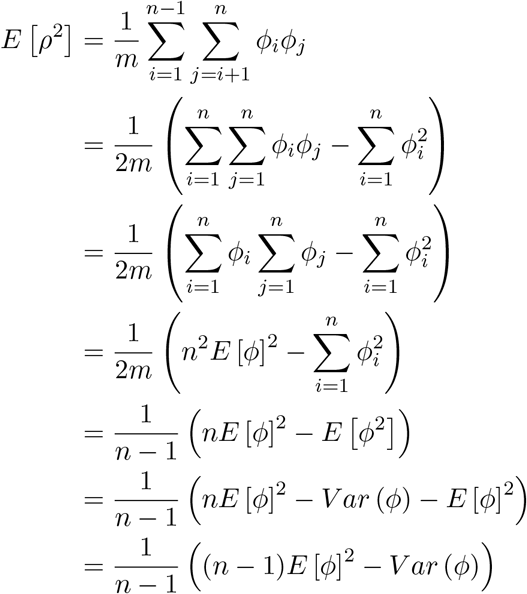

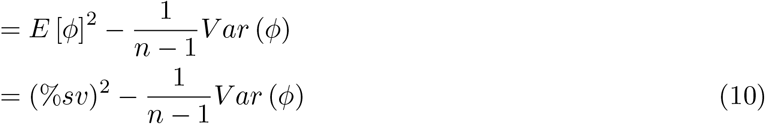

This provides an equation for the squared radius (i.e., squared distance from the origin) of a point in the *r*_sc_ mean versus *r*_sc_ s.d. plot. In the above derivation, *E* [*ϕ*] and *V ar* (*ϕ*) are taken across the percent shared variance of each neuron in the population *ϕ*_*i*_. Thus, *E* [*ϕ*] is equal to our population metric %sv. Now, we will bound *V ar* (*ϕ*), which by definition is greater than or equal to 0. Since 0 ≤ *ϕ*_*i*_ ≤ 1, one instance where the maximum variance occurs is when there are an equal number of *ϕ*_*i*_ = 0 and *ϕ*_*i*_ = 1 (and *E* [*ϕ*] = 0.5). Then,

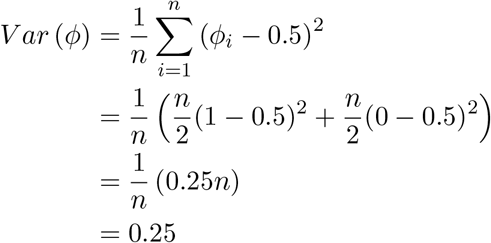

So 0 ≤ *V ar* (*ϕ*) ≤ 0.25. For a small number of neurons *n*, the second term is non-negligible. For example, for a model with 6 neurons and %sv = 50%, the radius of the data points may vary between 0.4472 and 0.5. As the number of neurons increases, the second terms becomes negligible, and data points lie approximately along an arc with radius equal to %sv. For example, for 30 neurons as in our simulations and a %sv of 50%, the radius only varies between 0.4913 and 0.5.

To summarize, equation (10) computes the distance from the origin of a point for a given population of neurons. For a fixed %sv, *V ar* (*ϕ*) can be the same or differ across many simulation runs. If *V ar* (*ϕ*) = 0 or is the same across runs, then the points will lie perfectly along an arc, with radius specified by equation (10). However, if *V ar* (*ϕ*) is different across runs, the distances of each point from the origin will differ slightly, so they will lie close to, but not exactly along, an arc.

With this analysis, we have shown that in the case of one latent dimensions:

- A point (i.e., corresponding to a given population of neurons, simulated or real) on the *r*_sc_ mean versus *r*_sc_ s.d. plot has distance from the origin (i.e., radius) less than or equal to %sv.
- If the %sv for individual neurons (*ϕ*_*i*_) are all the same (see Supplementary Math Note A), then the radius equals %sv.
- As the number of neurons increases, the radius becomes asymptotically closer to %sv.

##### C Relationship between correlation, loading similarity, and %sv (multiple latent dimensions)

In Supplementary Math Note A, we established a mathematical relationship between *r*_sc_, loading similarity, and %sv in the case of one latent dimension. Here, we generalize equation (8) to include multiple dimensions in order to better understand the relationship between *r*_sc_ and dimensionality. We demonstrate here that the general relationships between *r*_sc_, %sv, and loading similarity for one latent dimension also hold true for multiple latent dimensions. For multiple latent dimensions, the relative strengths of each dimension is an important consideration—a stronger dimension plays a bigger role in determining the *r*_sc_ distribution. Finally, we consider the relationship between dimensionality itself and *r*_sc_. We will discover below that increasing dimensionality tends to decrease the magnitude of *r*_sc_ values.

First, consider the case of two latent dimensions. Again, let *n* be the number of neurons, let **w** be the co-fluctuation pattern (i.e., loading vector [*w*_1_, *w*_2_, …, *w*_*n*_]^*T*^ ∈ ℝ^*n*×1^) with eigenvalue *λ*_*w*_, let **v** be another pattern orthogonal to **w** ([*υ*_1_, *υ*_2_, …, *υ*_*n*_]^*T*^ ∈ ℝ^*n*×1^; **v** ⊥ **w**), with eigenvalue *λ*_*υ*_, and let ψ ∈ ℝ^*n×n*^ be a diagonal matrix specifying the independent variance of each neuron (*ψ*_1_, *ψ*_2_, *…, ψ*_*n*_). Then the covariance is Σ = Σ_*shared*_ +ψ = Σ_*w*_ +Σ_*υ*_ +ψ = **w***λ*_*w*_**w**^*T*^ +**υ***λ*_*υ*_**v**^*T*^ +ψ. On the off-diagonals entries (i.e., if *i* ≠ *j*), Σ_*ij*_ = *λ*_*w*_*w*_*i*_*w*_*j*_ + *λ*_*υ*_*υ*_*i*_*υ*_*j*_. Along the diagonals, 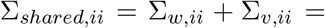 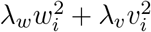 and 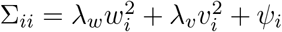.

Because the shared covariance matrix Σ_*shared*_ can be expressed as a sum of two component matrices Σ_*w*_ + Σ_*υ*_, we can express the %sv of neuron *i* (*ϕ*_*i*_) as

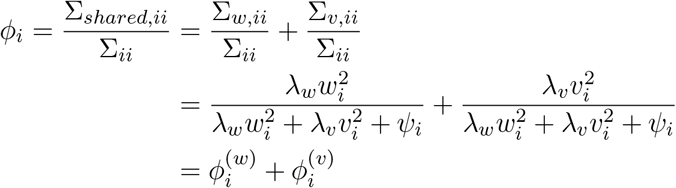

where 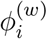 is the %sv variance of neuron *i* explained by dimension **w** and 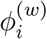 is the %sv variance of neuron *i* explained by dimension **v**.

With this decomposition of *ϕ*_*i*_, and following similar steps as in equation (8):

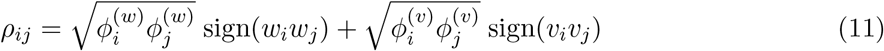

where %sv values (*ϕ*) are represented as proportions. Equation (11) relates *r*_sc_, %sv, and loading similarity for the case of two latent dimensions. Next, we compare these relationships for one versus two latent dimensions.

We will show that, for two latent dimensions, the relative strength of each dimension (i.e., the ratio *λ*_*w*_: *λ*_*υ*_) is an important consideration. For two latent dimensions, decreasing the overall %sv by decreasing both *ϕ*^(*w*)^ and *ϕ*^(*υ*)^ equally (e.g., *λ*_*w*_ = *λ*_*υ*_ and both decrease equally) pushes each *ρ*_*ij*_ closer to 0–*r*_sc_ mean and s.d. will decrease. This is similar to what happens for one latent dimension when %sv is decreased. On the other hand, even if the overall %sv is held constant, but *ϕ*^(*w*)^ increases relative to *ϕ*^(*υ*)^ (i.e., increase the strength of **w** relative to **v**), pairwise correlations could change. Each *ρ*_*ij*_ will largely be determined by *ϕ*^(*w*)^ and **w**—*r*_sc_ mean and s.d. will be more similar to what they would be if only **w** existed (Fig. 4*a*). In other words, each *ρ*_*ij*_ for two latent dimensions is the sum of the *ρ*_*ij*_ that would have been produced by each of the two constituent dimensions on their own. The dimension with larger relative strength *λ* will have larger *ϕ*; the stronger dimension will play a larger role in determining each value of *ρ*_*ij*_ and thus the resulting *r*_sc_ distribution.

Using this logic, we can deduce that increasing the loading similarity of one of the dimensions would increase *r*_sc_ mean and decrease *r*_sc_ s.d. for the same reasons as for one latent dimension (Supplementary Math Note A). Doing so for a relatively stronger dimension would result in larger changes in *r*_sc_ than doing so for a relatively weaker dimension.

We have shown how having multiple latent dimensions can affect the relationship between *r*_sc_, %sv, and loading similarity. Now, we show that dimensionality itself and *r*_sc_ are related—larger dimensionality tends to decrease *r*_sc_ mean and s.d. To see this, we can generalize equation (11) for *d < n* orthogonal latent dimensions **u**_**1**_, … , **u**_**d**_ ∈ ℝ^*n*^.

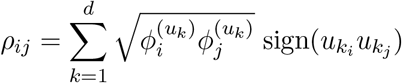

Considering the sign of one term, *ρ*_*ij*_ could have the same sign for 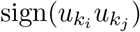 across all dimensions **u**_**1**_, … , **u**_**d**_; in this case, larger dimensionality acts to increase the correlation between neurons *i* and *j* (*ρ*_*ij*_) above the level corresponding to a single dimension. However, because the loading vectors **u**_**1**_, … , **u**_**d**_ are orthogonal, a pair of neurons *i* and *j* is likely to have many 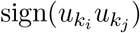 of opposite sign across dimensions; in this case, larger dimensionality pushes the correlation between neurons *i* and *j* (*ρ*_*ij*_) closer to 0. Thus, we would expect the magnitude of correlations to decrease as more dimensions are added (i.e., a tendency for *r*_sc_ mean and s.d. to decrease; Fig. 3*g*). In the next section, we show this relationship mathematically.

##### D Increasing dimensionality decreases arc radius

We establish here that increasing dimensionality results in a decrease in the radius of the arc in the *r*_sc_ mean versus *r*_sc_ s.d. plot (Fig. 3*g*). We extend the math for an arc for one latent dimension (Supplementary Math Note B) to multiple latent dimensions. We will refer to the one latent dimension as the ‘1-d case’ and multiple (*k*) latent dimensions as the ‘*k*-d case’.

We use the same notation as in Supplementary Math Note C. Consider the distance *c* of a point (corresponding to one instance of the population activity covariance matrix) from the origin in the *r*_sc_ mean versus *r*_sc_ s.d. plot. From Supplementary Math Note B, *c*^2^ = *E*[*ρ*^2^]. For this 2-d case, the correlation between neurons *i* and *j* is 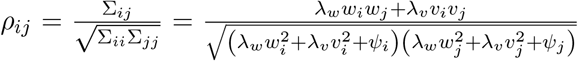.

Thus we can write 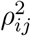 as:

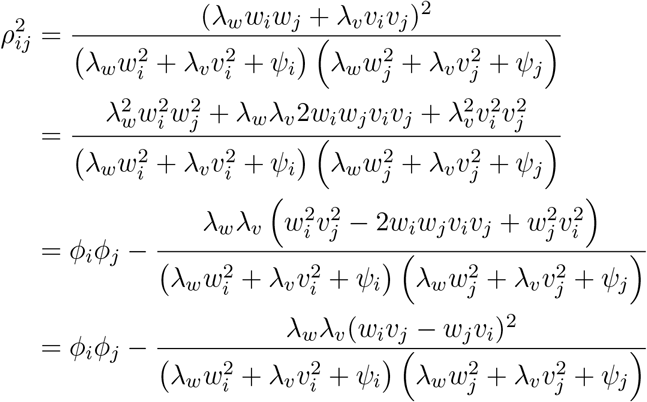

where the % shared variance of neuron *i* in this 2-d case is 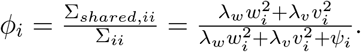.

Then letting *m* is the number of pairs in the population, and following similar steps to (10) in Supplementary Math Note B, we arrive at:

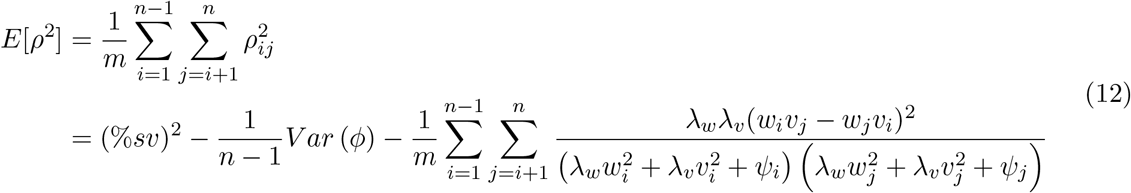

Not including the negative sign in front, note that this final term is non-negative (given that *λ*_*w*_ and *λ*_*υ*_ are non-negative, as for any covariance matrix). Thus, comparing the final line in equation (12) to the final line from equation (10), we observe that the distance of the point for the 2-d case in the *r*_sc_ mean versus *r*_sc_ s.d. plot is necessarily smaller than or equal to the distance for the corresponding 1-d case.

More generally, for a *k*-dimensional case we can show that:

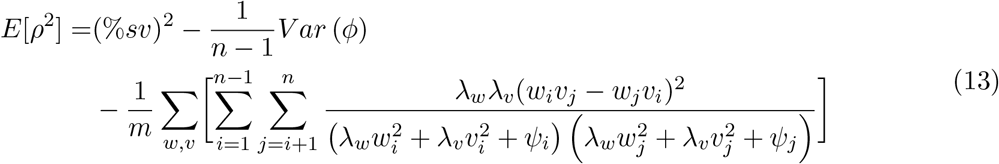

where the sum Σ_*w,v*_ is taken over all unique pairs of loading vectors (*w, v*). Indeed, as more latent dimensions are subsequently added, the radius of the *r*_sc_ mean versus *r*_sc_ s.d. plot decreases (Fig. 3*g*). Intuitively, this final term accounts for how population activity covaries along many different dimensions in the high-d firing rate space. As more *orthogonal* dimensions are added, population activity is further pulled in different directions in the high-d space, more interaction terms come into play, and the magnitude of correlations is further decreased. This tends to decrease both *r*_sc_ mean and *r*_sc_ s.d., explaining why the radius of the arc in the *r*_sc_ mean versus *r*_sc_ s.d. plot tends to decrease as dimensionality increases.

We note that *r*_sc_ mean and *r*_sc_ s.d. do not necessarily *both* need to decrease. For example, consider a pattern with a loading similarity of 1; loading weights for all neurons would have the same value, *r*_sc_ across all pairs would be the same value, and thus *r*_sc_ s.d. would be 0 (see Supplementary Math Note A). When a second pattern of necessarily low loading similarity (see Supplementary Math Note E) is added, *r*_sc_ values across pairs of neurons would differ, and *r*_sc_ s.d. would be larger than 0. Therefore, *r*_sc_ s.d. can increase when going from the 1-d case to the 2-d case. However, the corresponding decrease in *r*_sc_ mean would be larger in magnitude than the increase in *r*_sc_ s.d., resulting in an overall decrease in arc radius (Fig. 3*g*, 1 to 2 dimensions, data points closest to the horizontal axis).

The third term in equation (13) can also help explain variability of the radius (*E*[*ρ*^2^]) across different random instantiations with the same population metrics (Figs. 3*g* and 4). Consider a fixed %sv. For the 1-d case, the radius is determined by the first two terms of the above equation, and any variability in radius will be caused by different values of *V ar*(*ϕ*) across different instantiations. For the 2-d case, the third term also plays a factor in determining the radius, and this term varies across different random instantiations, typically to a larger degree than the second term for large numbers of neurons *n* (see Supplementary Math Note B). Thus, the 2-d and *k*-d cases have greater variability in *E*[*ρ*^2^] than 1-d cases (Fig. 3*g*, Fig. 4). Other subtle factors can affect the variability of *E*[*ρ*^2^]. For example, variability in *E*[*ρ*^2^] can increase or decrease depending on the relative strengths of each dimension and their corresponding loading similarities (Fig. 4 and Supplementary Fig. 5). This can be explained by the third component of equation (13), in particular by the terms involving *λ*_*w*_ and *λ*_*υ*_.

##### E Properties of loading similarities across different co-fluctuation patterns

We asked whether there was a relationship between the loading similarities of different co-fluctuation patterns in the same model. In our simulations and V4 data analysis, we ensured that we obtain unique co-fluctuation patterns by constraining dimensions to be orthogonal. Thus, we might conjecture that if one pattern has high loading similarity (e.g., [1, … , 1]), then another pattern in the same model necessarily has low loading similarity (e.g., [1, −1, 1, −1, … , −1, 1]). Indeed, this is true because the sum across the loading similarities of each pattern in a model is at most 1. We show this property of loading similarity here.

Let **w** and **v** be vectors representing two co-fluctuation patterns in the same model. We use the notation **w**· **v** to refer to the element-wise product between **w** and **v**, resulting in a vector that is the same size as **w** and **v**. Furthermore, we use *E*[**w**], *V ar*(**w**), and *Cov*(**w**) as shorthand to refer to computations across the elements of a vector (and *not* as operations on a random variable): e.g., 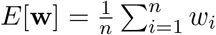, and 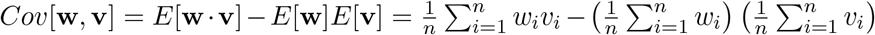. Also, in this section we refer to the loading similarity of vector **w** as *ls*(**w**) for shorthand.

We first show a constraint on loading similarities for a model with two co-fluctuation patterns (i.e. loading vectors for each dimension). Let *n* be the number of neurons and let **w**, **v** ∈ ℝ^*n*^ be two loading vecto As in our simulations and data analysis (see Methods), **w** and **v** are orthogonal unit vectors: 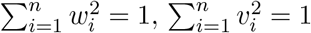, and 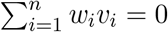. Then, using these constraints,

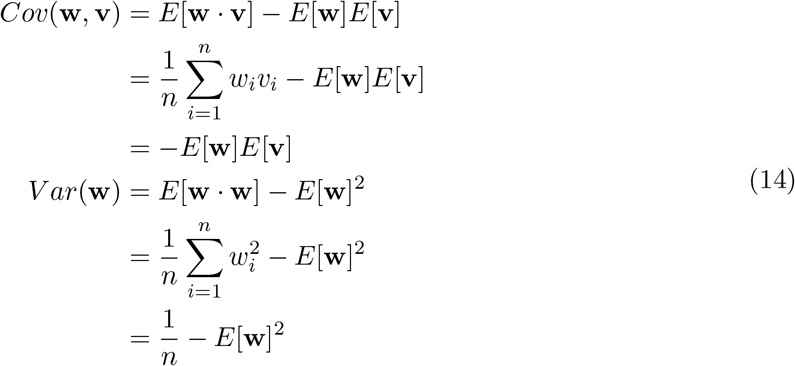

Because correlation is bounded between −1 and 1, we know that 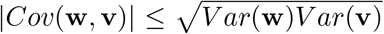. It follows that:

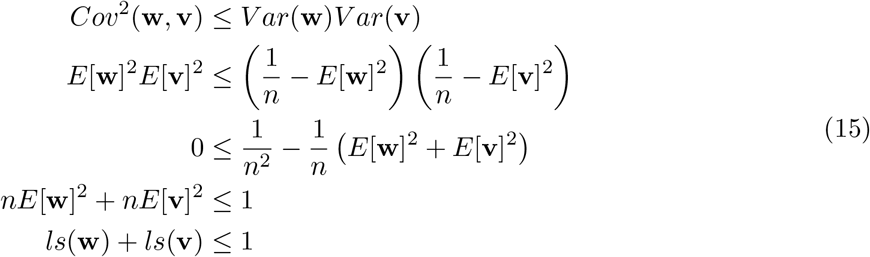

The last step follows from the definition of loading similarity:

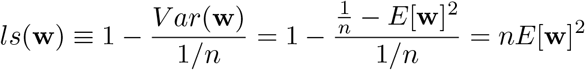

The final inequality in equation (15) proves the intuition provided at the beginning of this section–if *ls*(**w**) is large, then *ls*(**v**) must be small (at most 1 − *ls*(**w**)). More strongly, if *ls*(**w**) = 1, then *ls*(**v**) = 0.

Generally, for a model with *d* dimensions and patterns **u**_**1**_, … , **u**_**d**_ ∈ ℝ^*n*^, we can show that 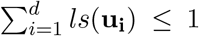. To see this, we can construct a matrix *C* with entries *c*_*ij*_ = *Cov*(**u**_**i**_, **u**_**j**_) = *j*, and 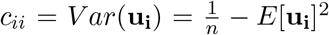 (derived from the constraints in equation (14)). Note that *C* ∈ ℝ^*d×d*^, with variances on the diagonal and covariances on off-diagonals, is a co-variance matrix, which implies *det*(*C*) ≥ 0. For a 3-d model, 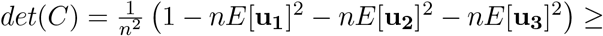, which implies *ls*(**u**_**1**_) + *ls*(**u**_**2**_) + *ls*(**u**_**3**_) ≤ 1. In general, for a *d*-dimensional model (with *d* ≤ *n*):

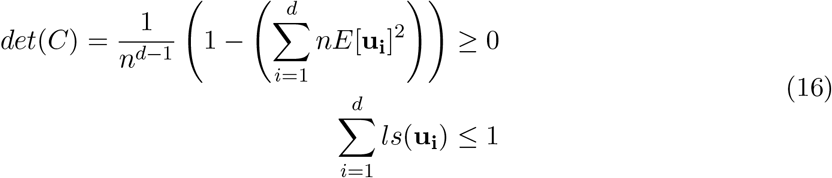

Equation (16) has several implications:

- If one knows the loading similarities of all dimensions **u**_**1**_, … , **u**_**d**_ in a model, then the maximum possible loading similarity of any new dimension is 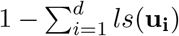. It follows that two dimensions with high loading similarity cannot co-exist in the same model.
- If one dimension has *ls* = 1, then all other dimensions in the model (or that would be added to the model) necessarily have *ls* = 0. Note that there is only one possibility for a pattern to have *ls* = 1 (i.e., 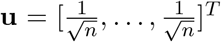, such that *Var*(**u**) = 0). This implies that there are many possibilities for a pattern to have *ls*(**u**) = 0. More loosely, there are relatively few ways for a pattern to have high loading similarity, but many more ways for a pattern to have low loading similarity.

##### F Maximum variance of a unit vector

We defined loading similarity for a co-fluctuation pattern **u** (normalized to have norm 1) of *n* neurons to be 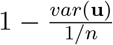, where the variance is computed along the elements of **u**. This value lies between 0 and 1 because the maximum variance across the elements of **u** is 1*/n*. We now show this mathematically.

Let **u** ∈ *R*^*n*^ be a unit vector. Because **u** is a unit vector, 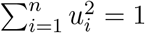. Using these facts:

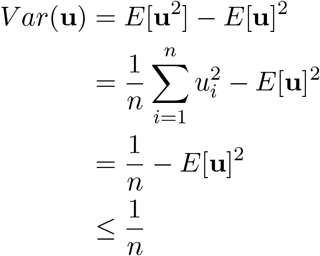

This holds with equality when *E*[**u**] = 0 (i.e., when the mean across the elements in a co-fluctuation pattern is 0). This implies that the smallest loading similarity is 0 (when *V ar*(**u**) = 1*/n*), and the largest loading similarity is 1 (when *V ar*(**u**) = 0).

## Supplementary Figures

**Supplementary Figure 1:**
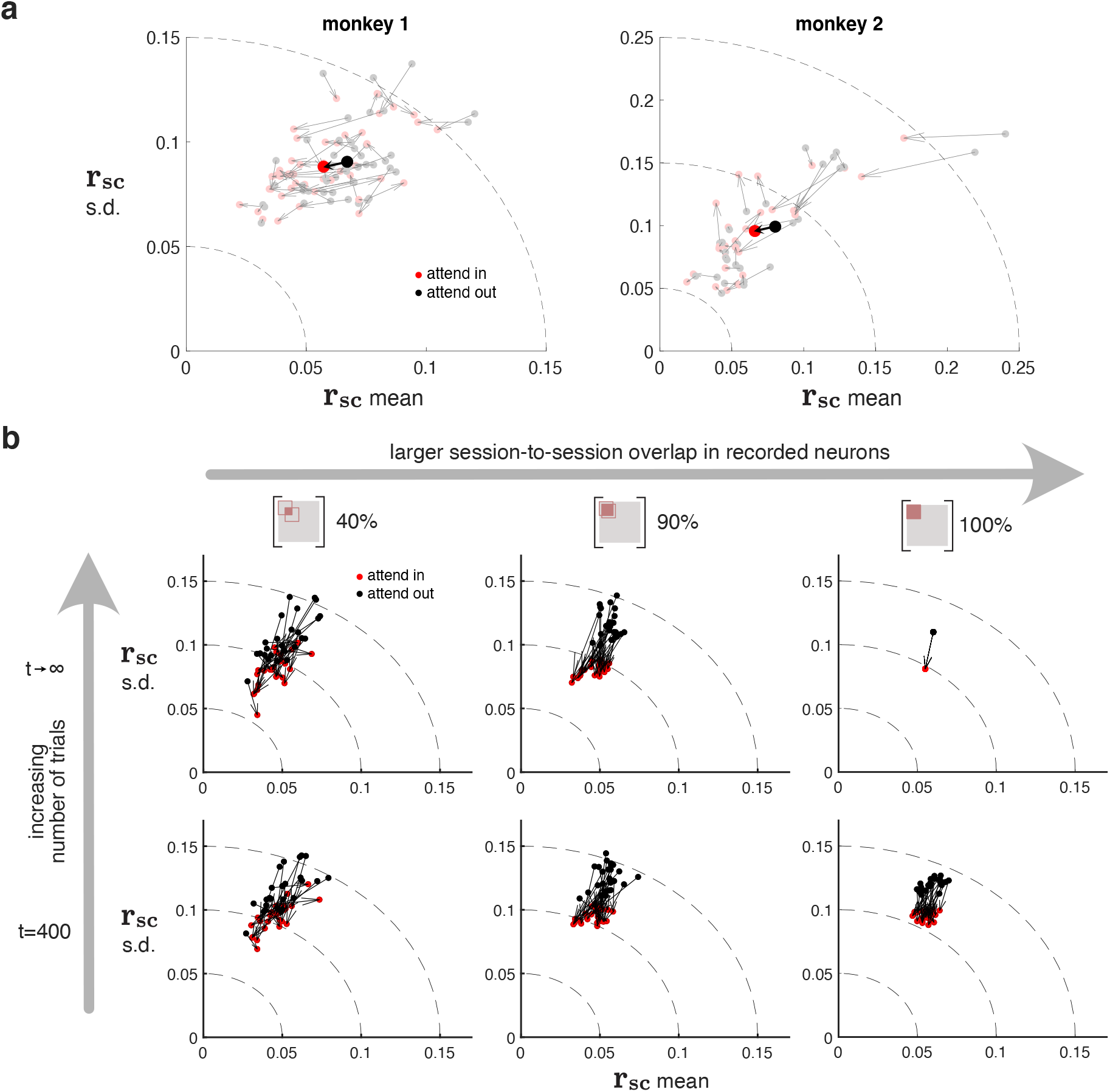
Session-to-session variability in pairwise metrics can be accounted for by finite trial counts and neuron sampling. We sought to understand to what extent the average pairwise correlations reported in Figure 6*b* varied from session to session, and the possible sources for this across-session variability. Here, we consider two possible sources: 1) the number of recorded trials, and 2) the overlap in recorded neurons across sessions. Although we likely recorded from many of the same neurons across sessions, the population of neurons recorded on each session was not identical due to recording instability (sometimes referred to as electrode drift), a well-known property of electrophysiological recordings. **a.** In V4 recordings, we observed substantial session-to-session variability in pairwise metrics. Each gray arrow, corresponding to an individual session, begins at *r*_sc_ mean and s.d. values for ‘attend-out’ (light gray dots) and ends at *r*_sc_ estimates for ‘attend-in’ (light red dots). The average change in *r*_sc_ mean and s.d (black arrow, black dot, solid red dot) correspond to the values we report in Figure 6*b*. **b.** Using simulations, we assessed how varying the number of trials and overlap of neurons across sessions contribute to the across-session variability of *r*_sc_ mean and s.d. The simulation procedure is described as follows. We first considered the setting of a large number of trials with varying amounts of neuron overlap. We created covariance matrix using factor analysis (equation 4) for 2,500 neurons and computed *r*_sc_ mean and s.d. from this matrix. Since we had access to the true covariance matrix, we considered this setting as having an “infinite” number of trials (top row). For each simulated recording session, we sampled 50 neurons from the set of 2,500 neurons. To mimic the effects of recording different neurons on different sessions, we sampled neurons such that a percentage of neurons were the same as those in the previous session (e.g., with 50 neurons in a session, a 40% overlap corresponds to 20 neurons being common with the previous session from two sessions, whereas 100% corresponds to having the same 50 neurons on all sessions). To simulate ‘attend in’ and ‘attend out’ trials, we chose FA parameters to be similar to those estimated from the V4 data (attend in: load. sim.=0.45, %sv=15%, dim.=3; attend out: load. sim.=0.5, %sv=20%, dim.=3). We found that as the percentage of neuron overlap between sessions increased, the across-session variability in the *r*_sc_ estimates decreased (moving left to right in top row). This indicates that if different neurons are recorded in different sessions, one should expect differences in estimates of pairwise metrics across sessions. We then considered how the number of recorded trials influences the across-session variability. We used the same simulation procedure as described above, except for one key difference. Instead of using the true covariance matrix to directly compute *r*_sc_, we instead simulated trials of “neuronal activity” from that covariance matrix. We simulated 400 trials because that was, on average, the number of trials we recorded per condition per session (i.e., each light-colored dot in panel *a* is computed from 400 trials). We then estimated the spike count covariance matrix based on these 400 trials. We found that the level of across-session variability did not substantially differ for 400 trials (bottom row) versus an infinite number of trials (top row), except in the case of 100% overlap, i.e. no recording instability. Using these simulations as a reference, it appears that much of the across-session variability in our reported V4 pairwise metrics (panel *a*) can be accounted for by neural recording instability and finite trial sampling. Additional across-session variability in the V4 results may be due to other cognitive factors, such as across-session differences in the animal’s motivation level.

**Supplementary Figure 2:**
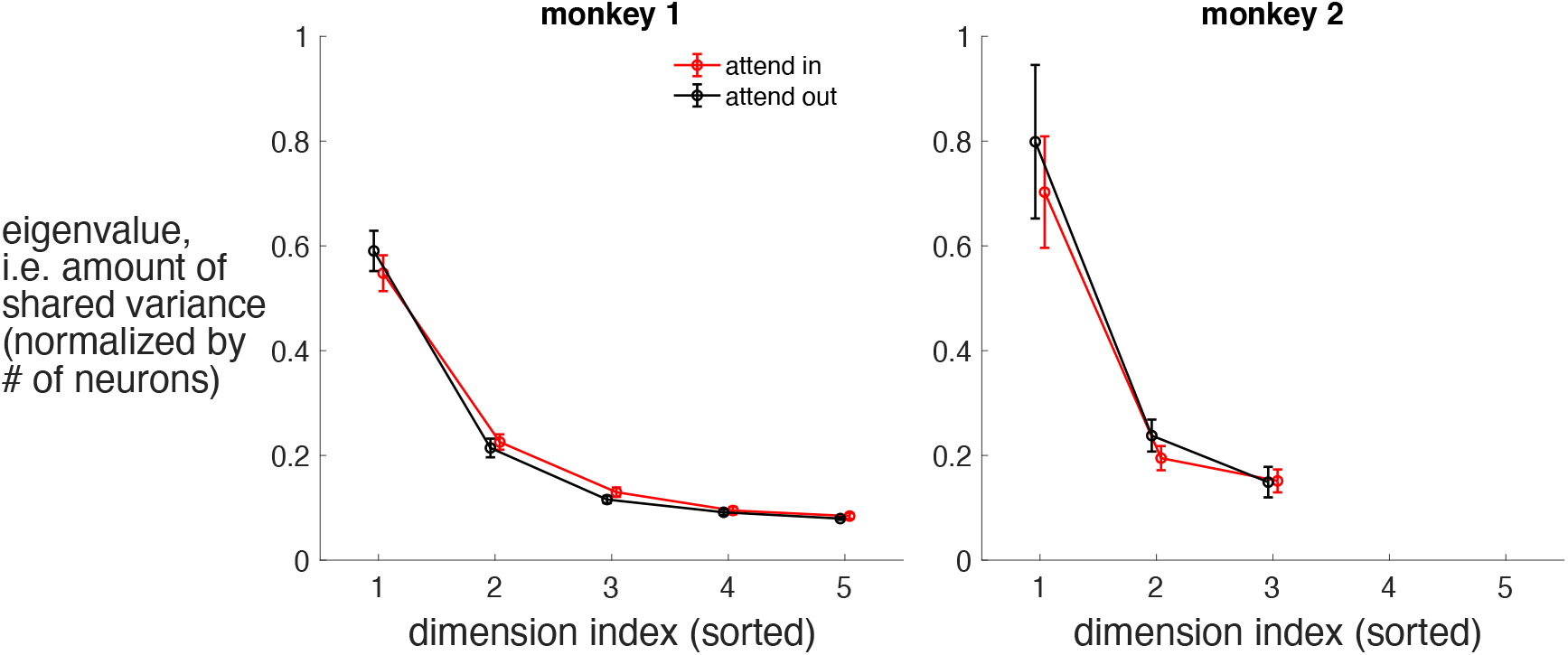
Eigenspectra for V4 population activity in ‘attend in’ and ‘attend out’ conditions. Although we observed only a modest change in dimensionality with attention (Fig. 6*c*), our simulations showed that the relative strength of each dimension (i.e., shape of the shared eigenspectrum) could alter the “effective dimensionality” of population activity and have large effects on pairwise metrics (Fig. 4*a*). Here, we asked whether the relative strengths of each dimension changed with attention. The eigenspectra were computed in the following way. We decomposed the V4 spike count covariance matrix into shared and independent components using factor analysis (see Methods). We then computed the eigendecomposition of the shared covariance matrix (Fig. 7, Σ_shared_ = *U* Λ*U*). We found that eigenvalues (diagonal of Λ) tended to increase linearly with the number of neurons recorded; therefore, in order to combine across sessions, we normalized the eigenvalues by dividing by the number of neurons recorded in each session. After normalizing, we computed the eigenspectrum averaged across sessions and stimulus orientations. Because the dimensionality identified by cross-validation differed across sessions, there were a different number of sessions that contributed to each average. We did not plot mean eigenvalues when there were fewer than 5 sessions to average (i.e., dimensions ≥ 6 for monkey 1; dimensions ≥ 4 for monkey 2). Error bars indicate standard error. Data points have been jittered horizontally for visual clarity. We found that the shape of the eigenspectra was qualitatively similar for ‘attend in’ and ‘attend out’ conditions (red and black curves have similar shape). In both conditions, the eigenvalues of the shared covariance matrix decayed (dot for each subsequent dimension was below dot for the previous dimension), indicating that a small number of dimensions were needed to explain the population-wide covariability. When comparing eigenspectra (i.e., the amount of shared variance explained by each dimension), one also needs to consider the firing rates under each condition. Mean firing rates tend to be higher for attend in than attend-out trials. Higher firing rates typically correspond to higher spike count variance due to the Poisson-like firing of neurons. All else being equal, the higher mean firing rates imply higher levels of both shared variance and independent variance (Churchland et al., 2010). Thus, a direct comparison of the eigenspectra should be done with caution. Nonetheless, we plotted attend-in and attend-out together to relate our results to previous reports (Huang et al., 2019; Ruff et al., 2019b). Consistent with these studies, we found that attention decreased the strength of the strongest dimension (red below black dot for dimension index 1), though the magnitude of the decrease we observed was more consistent with Ruff et al. (2019b) than Huang et al. (2019). Had we been able to equalize the mean firing rate across the two conditions, we likely would have observed an even greater difference between attend-in and attend-out. We note that the caveat described here for comparison of eigenspectra (i.e., the amount of shared variance) does not apply to comparisons of %sv (Fig. 6*c*) because %sv is normalized by the overall spike count variance.

**Supplementary Figure 3:**
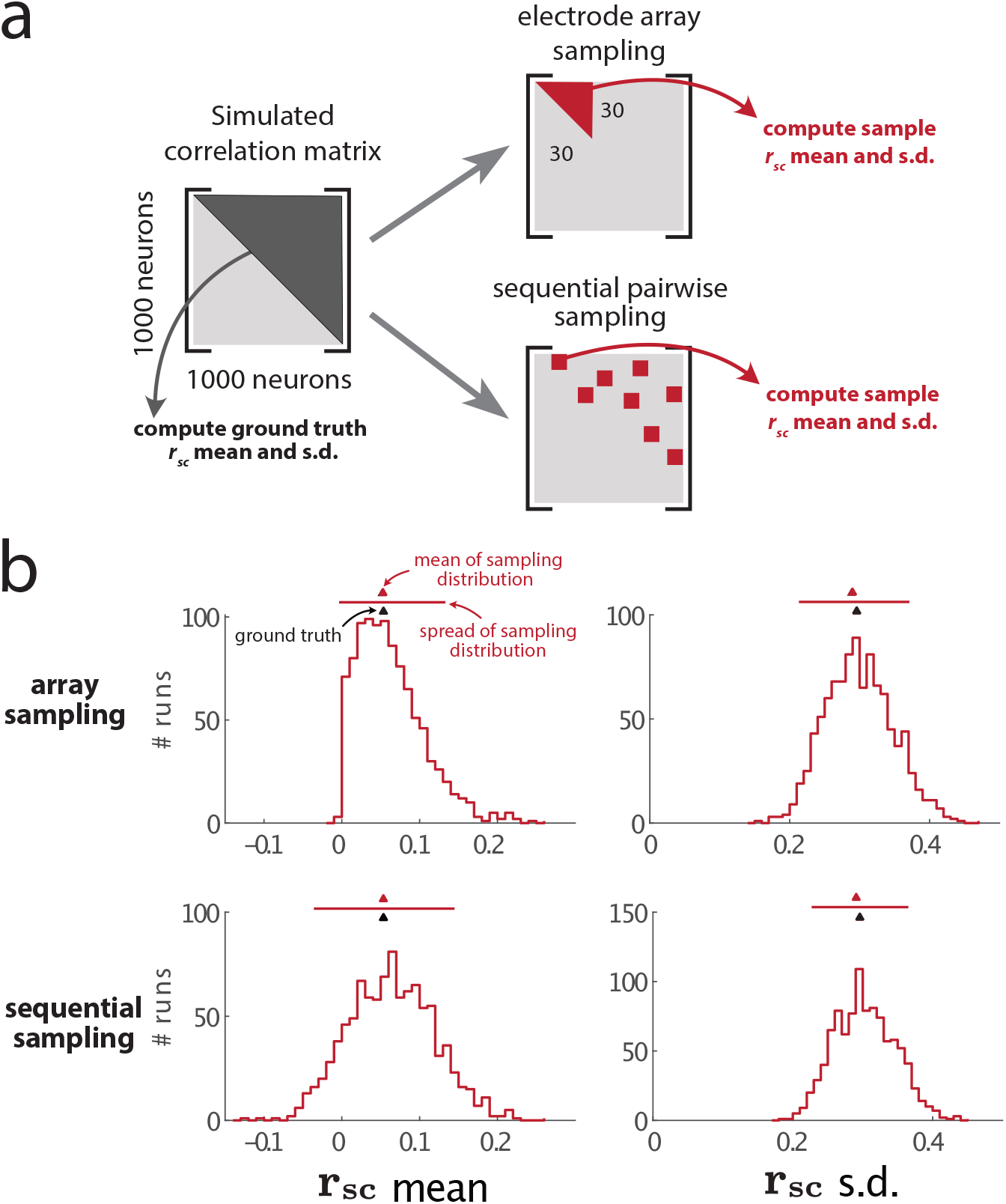
Pairwise metrics can be estimated equally well by either sampling pairs of neurons over many sessions or simultaneously sampling many neurons. Many studies that report *r*_sc_ mean measured the activity of one, or a few, pairs of neurons in each recording session, and combined *r*_sc_ across separate recording sessions. We wondered if our estimates of pairwise metrics may change if we could only sample random pairs of neurons, instead of simultaneously sampling a population of neurons using a Utah array (as in our V4 recordings). This “sequential pairwise sampling” is often done out of necessity due to physiological restrictions (e.g., recordings in deep brain structures) or technological limitations (e.g., recording with pairs of single recording electrodes, or a single tetrode). More recent studies often use multi-electrode array recording technologies, such as Utah arrays, that simultaneously record the activity of tens of neurons. We term this “electrode array sampling”; it is the type of sampling we considered in our main text simulations and V4 recordings. Here, we asked whether 1) sequential pairwise sampling and electrode array sampling provide similar estimates when computing pairwise statistics and 2) whether each type of sampling reflects the pairwise statistics of a larger neuronal population. **a.** To address these questions in simulation, we created, we created a large ‘ground truth’ covariance matrix of population activity, and then simulated recording sessions by sampling entries from this matrix using either a sequential pairwise or electrode array approach. We created a covariance matrix for a population of 1000 neurons using one co-fluctuation pattern (i.e., one latent dimension), a loading similarity of 0.2, and a %sv of 30% (Fig. 7). We also tried other values for the population metrics and obtained similar results to that shown below. From the covariance matrix, from which we computed a 1, 000 × 1, 000 correlation matrix (left, black upper triangle). The *r*_sc_ mean and s.d. of this correlation matrix were deemed to be the ground truth values that we aimed to estimate using sequential pairwise or electrode array sampling. To simulate an electrode array recording, we randomly sampled 30 neurons to obtain a 30 × 30 covariance matrix (top, red triangle). To simulate sequential recordings of pairs of neurons, we randomly sampled 30 entries of the population correlation matrix (bottom, random red squares). We repeated each type of sampling 1,000 times (i.e., 1,000 bootstrap runs), and for each run we computed *r*_sc_ mean and *r*_sc_ s.d. This procedure provided us with sampling distributions of the *r*_sc_ mean and s.d. **b.** For both array sampling (top row) and sequential pairwise sampling (bottom row), we compared the estimates of *r*_sc_ mean and s.d. on each run with the ground truth values. To assess whether either metric was consistently under or overestimated, we took the difference between the mean of the sampling distribution (red triangles) and ground truth values (black triangles). We also considered how consistent/replicable the estimates were, quantified by the spread of the sampling distribution (red bars, 90% confidence interval). For *r*_sc_ mean (left column), both array sampling (top row) and sequential sampling (bottom row) resulted in little to no statistical bias (red triangles and black triangles are closely aligned), and both types of sampling showed similar replicability (red bars are of similar lengths). We note that array sampling resulted in few negative *r*_sc_ mean estimates because the 30 × 30 correlation matrix obtained with array sampling must be a positive semi-definite matrix, limiting the range of possible *r*_sc_ mean values and making negative *r*_sc_ means unlikely. For *r*_sc_ s.d. (right column), both array sampling (top row) and sequential sampling (bottom row) slightly underestimated the ground truth value (red triangle lies to the left of black triangle). This underestimation bias was small relative to the larger variance in estimates of *r*_sc_ s.d. (red bars). Overall, we found that both sequential pairwise sampling and electrode array sampling provide similar answers for *r*_sc_ mean and *r*_sc_ s.d. Also, both types of sampling provide reasonably good estimates of *r*_sc_ mean and *r*_sc_ s.d. of a larger population, given a reasonable sampling of neurons (30 neurons for electrode, or 30 pairs for pairwise sampling here) relative to the larger population (1000 neurons here). This implies that it is still possible to gain insight into a larger population of neurons and use the intuitions provided in the main text of this work even in cases where one can only record sequentially from pairs of neurons over multiple recording sessions. Note that the results here assume the covariance matrix is accurately estimated using a large number of trials.

**Supplementary Figure 4:**
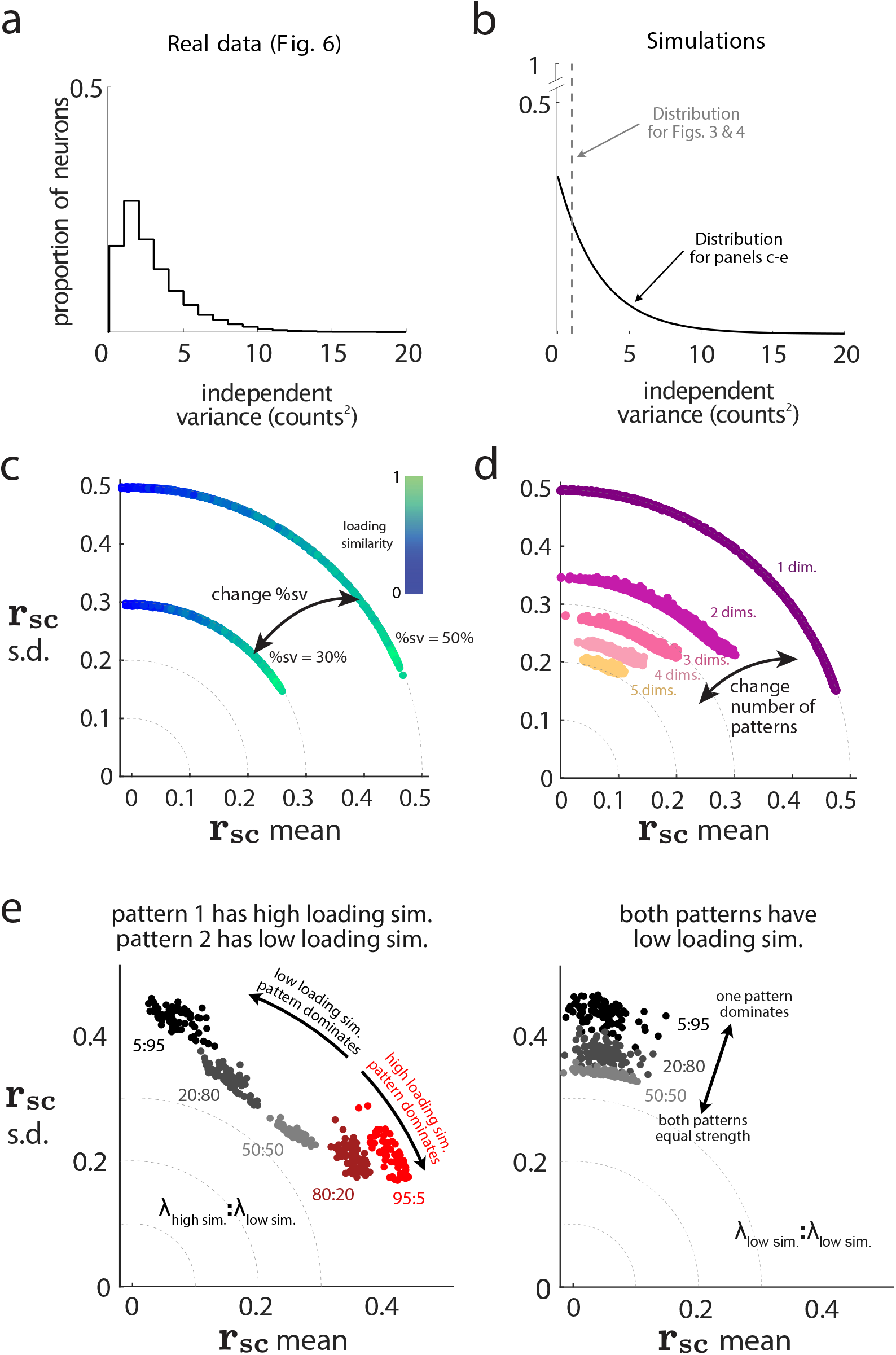
Relationship between pairwise and population metrics shows little dependence on distribution of independent variances. In our simulations (Figs. 3 and 4), we fixed the independent variance for each neuron to be the same value. However, we found that for real data, the independent variance differed across neurons. To account for this, we re-performed our simulations using a distribution of independent variances that was similar to that of the real data. We found the same trends, suggesting that different distributions of independent variances lead to similar conclusions about the relationship between pairwise and population metrics. **a.** We applied factor analysis to V4 population activity recorded during a spatial attention task (see Fig. 6 and Methods) and estimated each neuron’s independent variance (Fig. 7*a*, gray diagonal entries). Independent variances between neurons differed (dashed line denotes histogram). **b.** In the main text (Figs. 3 and 4), our simulations used a fixed independent variance of 1 for all neurons (dahsed gray line). Here (**c**-**e**), we re-performed the same simulations but drawing independent variances from an exponential distribution (solid black line) which approximates the distribution of the real data (compare solid black line here to that in in **a**). We used an exponential distribution with a mean of 3. **c.** Varying %sv and loading similarity while keeping the dimensionality fixed at 1. Same conventions as in Fig. 3*f*. We set %sv by scaling the eigenvalue of the one dimension (see Methods). With changing %sv, exponentially-distributed independent variances led to similar changes in radius of the arcs as when independent variances were all 1 (compare to Fig. 3*f*). However, here the arc lengths were smaller than in Figure 3*f* because small values of *r*_sc_ s.d. were unlikely with different independent variances. However, an *r*_sc_ s.d. of 0 is still possible under special conditions that are not likely to be encountered in practice (i.e., if the loading for each neuron is set to be the square root of its independent variance; see Supplementary Math Note A). **d.** Varying dimensionality and loading similarity while keeping %sv fixed at 50%. Same conventions as in Fig. 3*g*. We found that as dimensionality increases, *r*_sc_ mean and *r*_sc_ s.d. decrease, consistent with our simulations in Fig. 3*g*. **e.** Varying the relative strengths of two dimensions. Same conventions as in Fig. 4. We observed similar trends whether the independent variances were all fixed to 1 (Fig. 4) or drawn from an exponential distribution. However, here with exponentially-distributed independent variances, covariance matrices for which the pattern of the dominant dimension (i.e., the dimension with the largest eigenvalue) had a high loading similarity (left panel, 95:5, red dots) yielded a higher *r*_sc_ s.d. than when independent variances were the same for each neuron. This is for the same reasons as noted in panel **c**.

**Supplementary Figure 5:**
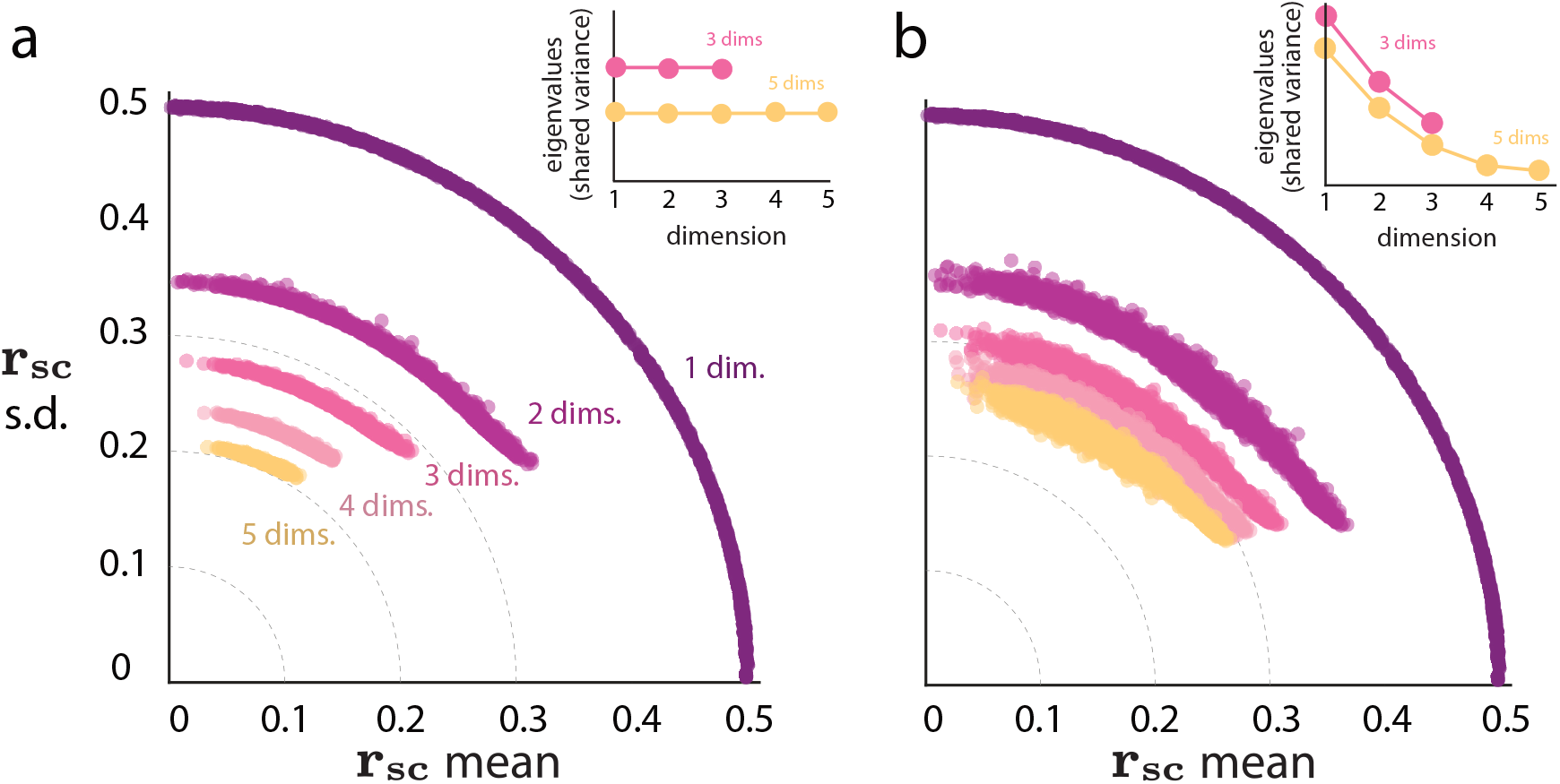
Relationship between pairwise and population metrics does not depend on relative strengths of dimensions. In Fig. 3*g*, we varied dimensionality and assessed the changes to pairwise metrics. For that simulation, we assumed that each dimension had equal strength (i.e., a flat eigenspectrum for the shared covariance matrix). Here, we asked if our simulation results held when we changed the shape of the shared covariance eigenspectrum such that it was no longer flat (i.e., some dimensions explained more shared variance than others). We found similar trends, suggesting that a different eigenspectrum shape leads to similar relationships between pairwise and population metrics as those found in the main text. **a.** Results for flat eigenspectra (reproduction of Fig. 3*g*, here with inset). Each dimension of the shared covariance matrix had the same eigenvalue (top right inset), and thus each dimension explained the same amount of shared variance. Note that models with different dimensionalities had different eigenvalues (inset, 3-dim. has larger eigenvalues than those for 5-dim.), because we scaled the eigenvalues (while keeping fixed each neuron’s independent variance) to enforce the %sv to be the same across models of different dimensionalities (inset, the sum of eigenvalues is equal for 3-dim. and 5-dim.). **b.** In neuronal recordings, the shape of eigenspectra curves estimated from recorded neuronal activity is usually not flat (see Supplementary Fig. 2). We sought to understand how the shape of the eigenspectrum affects the relationship between pairwise and population metrics. We re-performed the simulations in Figure 3*g* except here with the eigenspectra curves were exponentially decaying according to 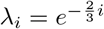 (inset). The overall trend held true: an increase in dimensionality led to an overall decrease in the radius of the arc (yellow dots closer to origin than purple dots). We also observed two subtle differences from Figure 3*g*. First, an exponentially-decaying eigenspectrum tended to have a higher *r*_sc_ mean and *r*_sc_ s.d. compared to its corresponding flat eigenspectrum (e.g., yellow dots for 5 dims are further from origin here than in panel *a*). This occurs because, for an exponentially-decaying eigenspectrum, an added dimension explains relatively little shared variance (right side of the curves in inset). Thus, the added dimension results in only small changes in *r*_sc_ mean and *r*_sc_ s.d. On the other hand, adding a dimension to the flat eigenspectrum affects *r*_sc_ mean and *r*_sc_ s.d. as much as any other dimension, leading to larger changes in *r*_sc_ mean and *r*_sc_ s.d. than in the case of an exponentially-decaying eigenspectrum. The second difference was, for a given dimensionality, a greater radial and angular spread for exponentially-decaying eigenspectra compared to flat eigenspectra (e.g., yellow dots for 5 dims are more spread out in the radial direction here than in panel *a*; yellow dots are also more spread out parallel to the dashed lines). This occurs because, when the eigenspectra are not flat, there is greater diversity in how the co-fluctuation patterns of different dimensions can contribute to *r*_sc_. In other words, switching the eigenvalues of two dimensions with equal eigenvalues (i.e., both dimensions explain the same amount of shared variance) results in the same model and same covariance matrix—yielding the same values for *r*_sc_ mean and *r*_sc_ s.d. However, switching the strengths of two dimensions with different eigenvalues is likely to result in a different covariance matrix and thus different values of *r*_sc_ mean and *r*_sc_ s.d. Thus, for non-flat eigenspectra, the greater diversity by which co-fluctuation patterns can contribute to the shared covariance matrix leads to greater spread in the *r*_sc_ s.d. vs *r*_sc_ mean plots. The mathematical details about the spread of dots in the radial direction are provided in Supplementary Math Note D. An implication of this analysis is that it is important to report the eigenspectrum shape whenever one reports dimensionality. For example, consider population activity described by 5 dimensions with an exponentially-decaying eigenspectrum (panel *b*, yellow dots). Because of 1) low shared variance in the last 2 dimensions, and 2) greater diversity in how dimensions contribute to the shared covariance matrix, the activity might closely resemble that described by 3 dimensions with a flat eigenspectrum (*a*, pink dots). Thus, considering both dimensionality and the eigenspectrum curve, instead of dimensionality alone, will lead to a more complete picture of the structure of population activity.

## Notes

### Competing Interest Statement

The authors have declared no competing interest.

